# Negative durotaxis: cell movement toward softer environments

**DOI:** 10.1101/2020.10.27.357178

**Authors:** Aleksi Isomursu, Keun-Young Park, Jay Hou, Bo Cheng, Ghaidan Shamsan, Benjamin Fuller, Jesse Kasim, M. Mohsen Mahmoodi, Tian Jian Lu, Guy M. Genin, Feng Xu, Min Lin, Mark Distefano, Johanna Ivaska, David J. Odde

## Abstract

Durotaxis – the ability of cells to sense and migrate along stiffness gradients – is important for embryonic development and has been implicated in pathologies including fibrosis and cancer. Although cellular processes can sometimes turn toward softer environments, durotaxis at the level of cells has thus far been observed exclusively as migration from soft to stiff regions. The molecular basis of durotaxis, especially the factors that contribute to different durotactic behaviors in various cell types, are still inadequately understood. With the recent discovery of ‘optimal stiffness’, where cells generate maximal traction forces on substrates in an intermediate stiffness range, we hypothesized that some migratory cells may be capable of moving away from stiff environments and toward matrix on which they can generate more traction. Combining hydrogel-based stiffness gradients, live-cell imaging, genetic manipulations, and computational modeling, we found that cells move preferentially toward their stiffness optimum for maximal force transmission. Importantly, we directly observed biased migration toward softer environments, i.e. ‘negative durotaxis’, in human glioblastoma cells. This directional migration did not coincide with changes in FAK, ERK or YAP signaling, or with altered actomyosin contractility. Instead, integrin-mediated adhesion and motor-clutch dynamics alone are sufficient to generate asymmetric traction to drive both positive and negative durotaxis. We verified this mechanistically by applying a motor-clutch-based model to explain negative durotaxis in the glioblastoma cells and in neurites, and experimentally by switching breast cancer cells from positive to negative durotaxis via talin downregulation. Our results identify the likely molecular mechanisms of durotaxis, with a cell’s contractile and adhesive machinery dictating its capacity to exert traction on mechanically distinct substrates, directing cell migration.

---

The capacity of living cells to undergo controlled migration is critical for tissue homeostasis and development, and underlies pathological conditions like cancer metastasis (*1*, *2*). Cells migrate in response to chemical and physical cues including the elasticity, or stiffness, of the surrounding extracellular matrix (ECM). The well-known tendency for many cells to migrate toward stiffer substrates, known as durotaxis (*3*–*8*), has implications for both developmental morphogenesis (*9*, *10*) and cancer cell invasion (*8*, *11*).

Despite progress in empirically identifying environmental conditions and molecular components that enable or promote durotaxis (*4*, *5*, *12*–*14*), our understanding of its fundamental mechanisms in different cell types is lacking. A long-standing mathematical model for cell migration is based on the motor-clutch mechanism (*15*–*18*), in which F-actin filaments polymerize against the plasma membrane to push the cell edge forward, while being simultaneously pulled away from the cell edge by ATP-dependent myosin II (‘molecular motors’) and pushed by force from the ATP-dependent polymerization itself. Retrograde F-actin flow can be mitigated by mechanical connections or ‘clutches’, typically integrin-mediated adhesions, between the F-actin and ECM to generate traction and bias cell movement toward more adhesive environments (*19*, *20*). Similarly, fibroblasts on stiffness gradients exhibit asymmetric traction, which has been postulated to contribute to their polarization and durotaxis (*6*, *21*). Interactions between actomyosin machinery and integrin-mediated adhesions have also been implicated in neuronal growth and pathfinding; however, the unifying principles underlying these behaviors across cell types have not been established (*22*–*24*).

Recently, cellular traction forces were shown to be maximal on substrates of an ‘optimal stiffness’ that can be predicted by the motor-clutch model (*17*, *18*, *25*–*29*). However, the biological relevance of this on cell behavior remains to be fully elucidated. Due to the key role of traction in driving mesenchymal cell migration, we predicted that any cell whose adhesion dynamics are governed by the motor-clutch model could potentially migrate toward *softer* environments, if such environments were closer to the cell’s optimal stiffness for maximal traction generation. We call this behavior ‘negative durotaxis’.

To test our hypothesis, we seeded U-251MG human glioblastoma cells, previously shown to exhibit maximal traction at an optimal stiffness of 5‒10 kPa (Fig. 1a)(*28*), on fibronectin-functionalized polyacrylamide hydrogels having a continuous stiffness gradient of approximately 0.5‒22 kPa (Fig. 1b)(*30*) – a range representative of healthy and malignant brain tissue (*31*). We observed a strong tendency for these cells to undergo negative durotaxis, migrating from the stiffest regions to regions of intermediate stiffness over time (Fig. 1b‒c). Fewer cells were observed in the softest regions, implying that cells below the optimal stiffness underwent conventional positive durotaxis. To exclude cell proliferation as a cause of these differences, we quantified the rate of EdU incorporation in cells cultured on homogeneous 0.5, 9.6 and 60 kPa substrates. Proliferation was equal on 9.6 kPa and 60 kPa hydrogels and only slightly lower on 0.5 kPa substrates (Fig. S1a‒b), suggesting that the absence of cells in the stiffer regions of the gradient was indeed due to biased migration.

**Figure 1.**
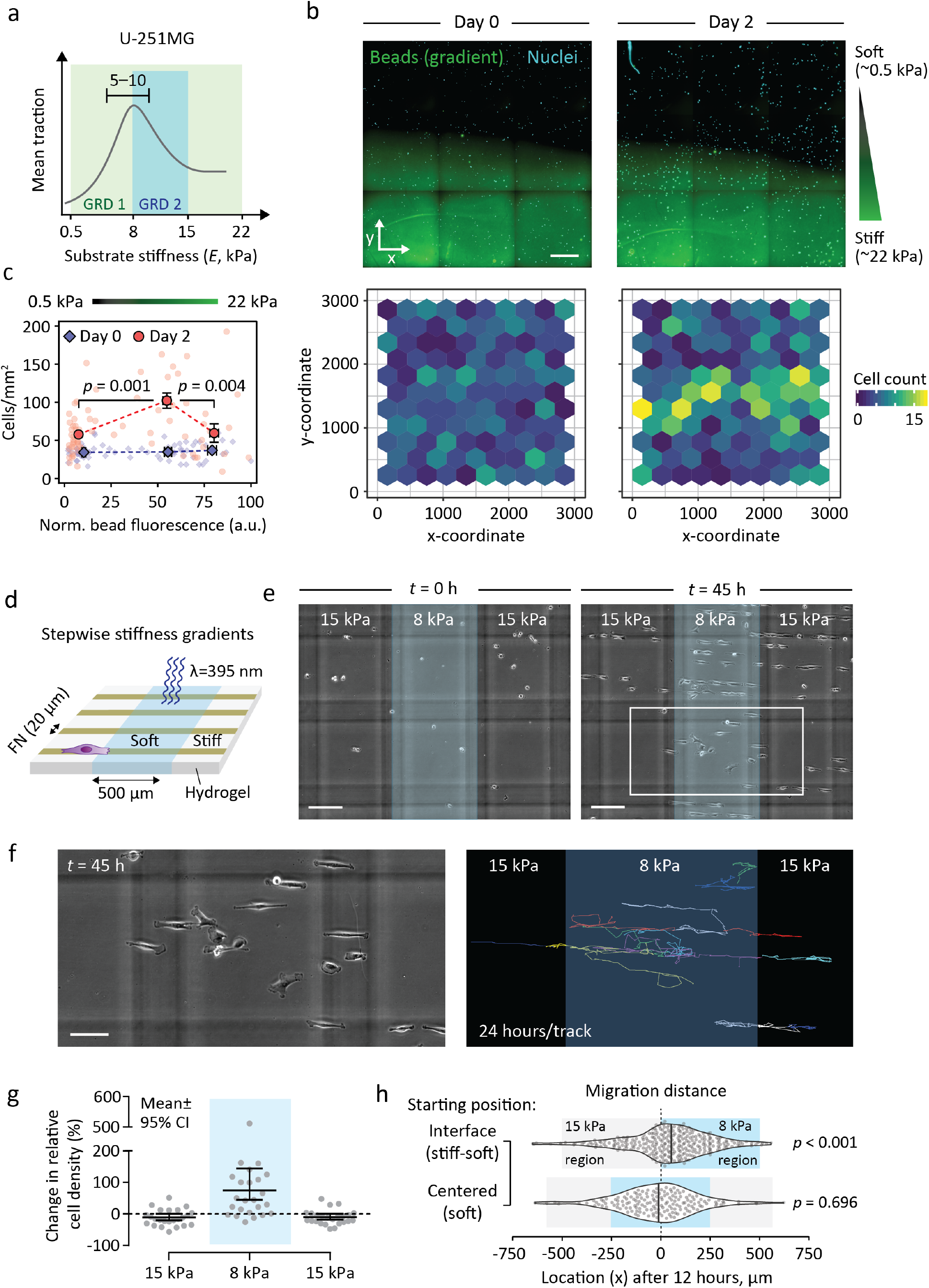
U-251MG glioblastoma cells undergo negative durotaxis. (**a**) Schematic representation of U-251MG traction, maximal on 5‒10 kPa substrates (*28*), and how it relates to the two stiffness gradients employed here. (**b**) (Top) Representative region of a diffusion-based polyacrylamide stiffness gradient (Young’s modulus ~0.5–22 kPa), at the outset of the experiment and 48 hours later. U-251MG cells are indicated by nuclear staining. Scale bar, 500 μm. (Bottom) Quantification of cells across the gradient. (**c**) Cell density in different parts of the stiffness gradient. Bins denote pooled regions of interest in the bottom, middle and top third of the gradient, respectively. Mean ± SEM of n = 14‒42 ROIs, analyzed by Kruskal-Wallis one-way ANOVA and Dunn’s *post hoc* test. (**d**) Schematic representation of photoresponsive hydrogels with stepwise stiffness gradients. (**e–h**) U-251MG migration on stepwise gradient hydrogels. A representative example (e) and quantification (g) of the change in cell density across the gradients over time. Blue overlay denotes softer, UV-exposed regions. Vertical and horizontal gray lines in (e) are out-of-focus markings in the underlying glass, used as a reference. Scale bar, 200 μm. Mean ± 95% CI from n = 24 fields of view, from two independent experiments. (f) End points (left) and 24-hour tracks (right) depicting the migration of individual cells in the region denoted by a white rectangle in (e). Scale bar, 100 μm. (h) Violin plots of accumulated distance migrated by individual cells along the x-axis over 12 hours, starting from a gradient (top) or from the middle of a compliant region (bottom). Vertical lines denote medians, n = 164‒296 cells from two independent experiments. Analyzed by sign test.

To verify this, we cultured cells on photoresponsive hydrogels with alternating 8 and 15 kPa regions, connected by steep stiffness gradients (hereafter ‘stepwise gradients’) (Figs. S2, S3; Supplementary Text 1). 20 μm wide fibronectin lines were printed across the gradients to facilitate cell motility. Live imaging revealed that cells migrated along the fibronectin lines and clustered preferentially in the softer 8 kPa regions (Figs. 1d–e, g; S1c–d; Movie S1). Moreover, tracking of individual U-251MGs confirmed that any cells making contact with a stiffness gradient migrated preferentially to the 8 kPa side (Fig. 1f, h; Movie S2). Taken together, these data demonstrate that U-251MGs are capable of negative durotaxis from stiff to soft environments, consistent with their stiffness optimum for maximal traction.

To gain insight into the molecular basis of negative durotaxis, we investigated key mediators of mechanotransduction, whereby biomechanical cues are translated into changes in cell signaling and behavior (*32*). We speculated that a biphasic response in any of these could, in part, modulate the negative durotaxis of U-251MGs. However, no changes were observed in myosin II light chain (MLC2), focal adhesion kinase (FAK) or extracellular signal-regulated kinase (ERK) phosphorylation in U-251MGs cultured on substrates with moduli of 0.5, 8 or 50 kPa (Fig. 2a‒b). These results were surprising because, in most adherent cell types, increasing substrate stiffness supports integrin clustering and focal adhesion (FA) growth, promoting the activation of mechanosensitive downstream signaling pathways (*18*, *33*, *34*).

**Figure 2.**
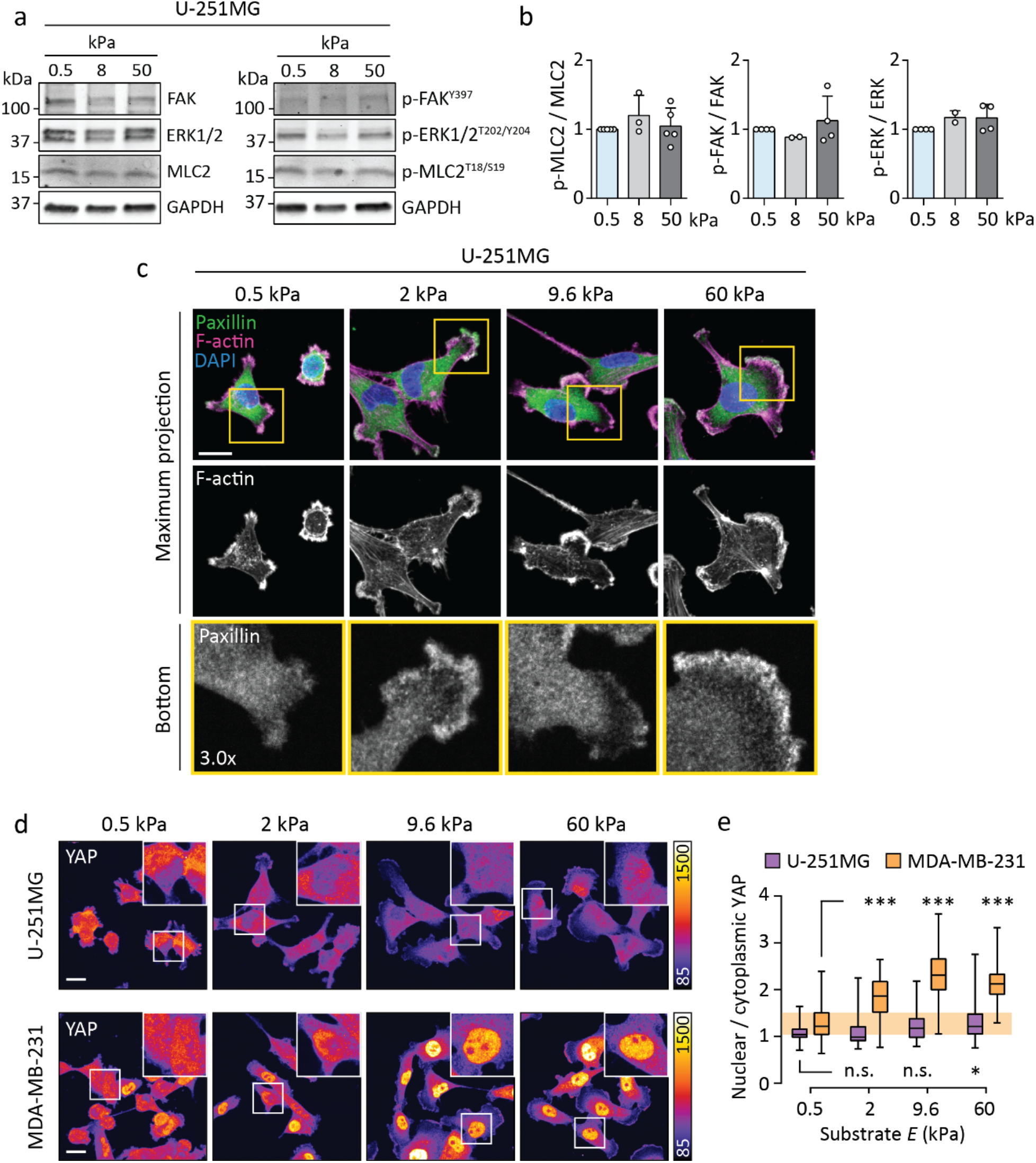
U-251MG cells display limited mechanosensitive signaling and adhesion maturation. (**a–b**) Representative western blot (a) and quantification (b) depicting protein phosphorylation in U-251MGs on 0.5‒50 kPa substrates. Mean ± SD of 2‒5 independent experiments. (**c**) Immunofluorescence images of paxillin and F-actin in U-251MGs on 0.5‒60 kPa substrates. The bottom panels show individual focal planes from confocal stacks, corresponding to the basal side of each cell. Scale bar, 20 μm. (**d–e**) Immunofluorescence images (d) and quantification (e) showing the intracellular localization of YAP as a function of substrate stiffness in U-251MG and MDA-MB-231 cells. Insets depict representative nuclei. Scale bar, 20 μm. Each box displays upper and lower quartiles and a median, the whiskers denote minimum and maximum values. n = 57‒ 135 cells, ***p < 0.001, *p = 0.018, n.s. = not significant, analyzed by Kruskal-Wallis one-way ANOVA and Dunn’s *post hoc* test.

This prompted us to compare focal adhesions in U-251MGs, capable of negative durotaxis, and MDA-MB-231 breast adenocarcinoma cells, which reportedly undergo positive durotaxis (*8*). As expected, MDA-MB-231s displayed stiffness-induced growth of paxillin-positive FAs (Fig. S4a) whereas U-251MGs displayed very few FAs even on 60 kPa substrates, as confirmed by immunostaining of paxillin (Fig. 2c) and additional FA markers, vinculin and phosphorylated FAK (Fig. S4b). This was not due to low expression of mechanosensitive adhesion proteins talin-1, talin-2 or vinculin, or due to low myosin II activity (p-MLC2), as these were expressed at comparable levels in U-251MG, MDA-MB-231, and human osteosarcoma U-2 OS, another FA-forming (*35*) cell line (Fig. S4c‒d). Nevertheless, U-251MGs displayed high β1-integrin activity and their spreading on fibronectin was sensitive to β1-integrin inhibition with a function-blocking antibody (Mab13) (Fig. S4e‒g), suggesting that they interact with their substrate primarily through integrins.

Hippo-family proteins yes-associated protein 1 (YAP) and transcriptional co-activator with PDZ-binding motif (TAZ) are transcriptional co-regulators that integrate cues from different mechanical and biochemical sources to direct cell behavior. Nuclear localization and activation of YAP/TAZ on stiff substrates are linked to increased F-actin assembly and FA formation; conversely, YAP/TAZ can promote adhesion turnover and cell migration (*36*) and baseline YAP activity may even be necessary for conventional durotaxis (*13*). We stained endogenous YAP from MDA-MB-231s and observed robust stiffness-induced nuclear translocation (Fig. 2d–e). In contrast, U-251MGs displayed much lower nuclear YAP on both soft and stiff substrates, with a slight increase but no visible peak between 0.5 and 60 kPa (Fig. 2d–e). Thus, mechanosensitive signaling responses of U-251MGs are minimal and not specific to the 5‒10 kPa range, and cannot explain negative durotaxis.

The optimal stiffness for U-251MG traction and the increasing overall motility of these cells (random motility coefficient, RMC) with stiffness up to 100 kPa can be explained by motor-clutch dynamics (*28*). Without talin unfolding and vinculin-mediated ‘clutch reinforcement’ and FA growth, the motor-clutch model naturally predicts a biphasic dependence of traction forces on substrate stiffness (*18*). After confirming that U-251MGs migrated preferentially toward their known traction optimum in all of our experimental conditions (Fig. 1a‒h), we investigated whether stochastic computational simulation of cell-level motor-clutch dynamics would be sufficient to reproduce negative durotaxis (Fig. 3a, Supplementary Text 2). We simulated the migration of individual U-251MGs on mechanically homogeneous substrates for one hour to allow the system to reach a dynamic steady state, then placed each cell on a continuous substrate consisting of alternating 60 μm wide regions of low and high stiffness, joined together by smooth 30 μm wide stiffness gradients (Figs. 3b, S5a‒b).

**Figure 3.**
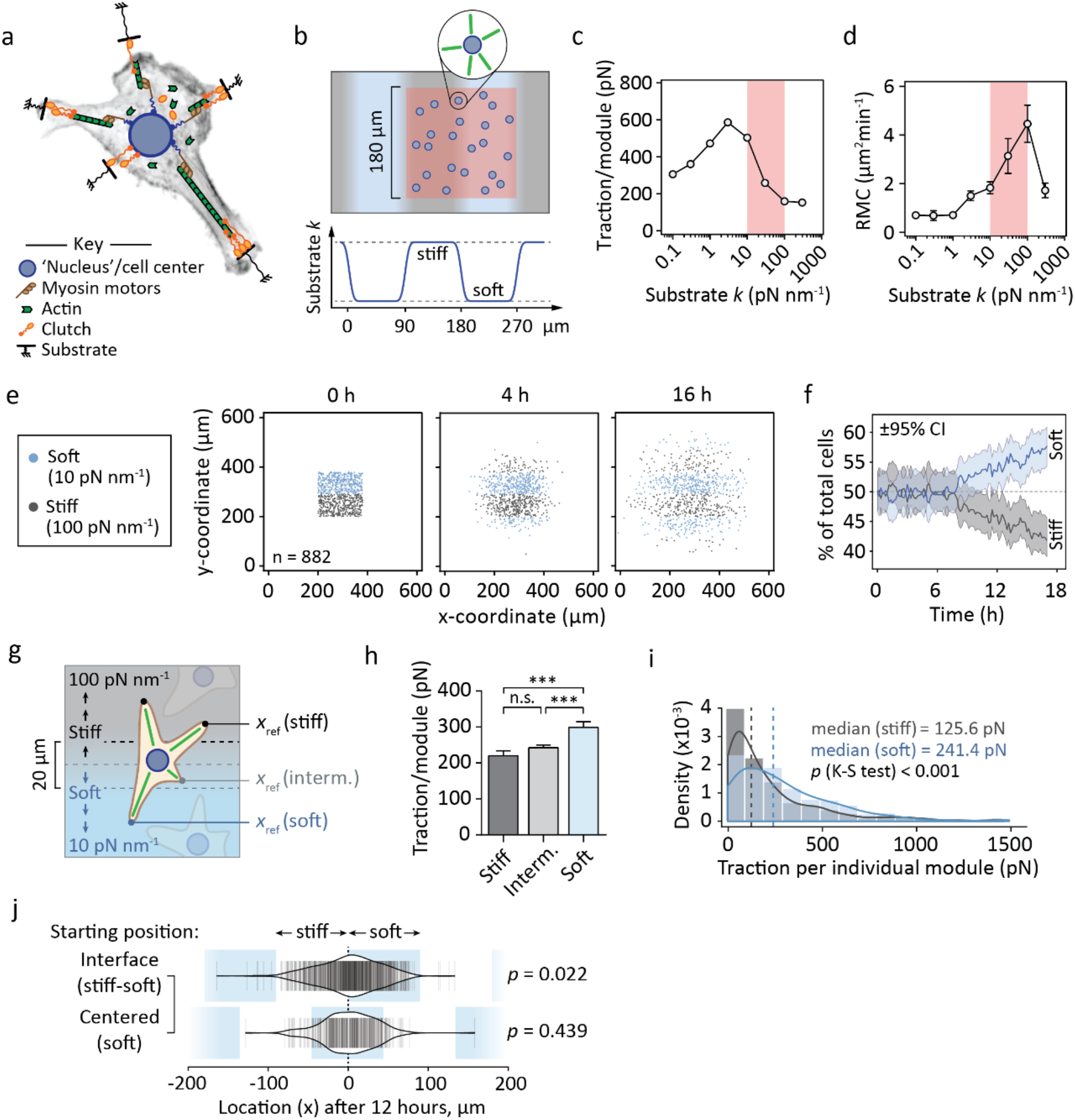
Motor-clutch simulations recapitulate negative durotaxis. (**a**) Schematic representation of the cell migration simulator (*28*). Individual modules and a central cell body are attached to the elastic substrate by sets of clutch molecules (Supplementary Text 2). (**b**) Experimental setup used here and in (Figs. S6). Simulated cells in a dynamic steady state were placed on a substrate with repeating stiff and soft regions and tracked over time. An equal number of cells were placed on both stiffnesses (red area). (**c‒d**) Module-wise traction forces (c) and RMC (d) of the simulated cells as a function of substrate stiffness. Red overlay highlights the range of the 10‒100 pN nm^−1^ gradient in (e‒j). Mean ± SEM of n = 10 cells. (**e–f**) Evolution of cell density on mechanically heterogeneous substrates over time. (e) Coordinates of individual cells 0, 4 and 16 hours into the simulation. Stiff (≥55 pN nm^−1^) and compliant (<55 pN nm^−1^) regions are indicated by gray and blue, respectively. (f) Fraction of cells residing in the stiff and soft regions over the course of the simulation. ±95% CI, n = 882 cells. (**g**) When individual cells were on top of a stiffness gradient, their traction forces were recorded. (**h**–**i**) Forces exerted by clutch modules on stiff, intermediate and soft substrate, while the cell is on top of a stiffness gradient. (h) Bar graphs depicting mean ± SEM of n = 292‒1380 modules. ***p < 0.001, n.s. = not significant, Kruskal-Wallis one-way ANOVA and Dunn’s *post hoc* test. (i) Histograms overlaid with probability density functions, dashed lines indicate medians. n = 292‒365 modules, analyzed by Kolmogorov-Smirnov test. (**j**) Violin plots of accumulated distance migrated by individual cells along the orientation of the gradient and over 12 hours, starting from a gradient (top) or from the middle of a compliant region (bottom). n = 326‒759 cells, analyzed by sign test.

On 10‒100 pN nm^−1^ gradients, corresponding to ~10‒100 kPa for typical adhesion sizes (*37*), and where the cells’ optimal stiffness overlaps with the softer regions (Fig. 3c‒d), we found that the majority of cells translocated away from stiffer areas in the first 12 hours of the simulation (Fig. 3e‒f). This occurred despite the cells being less motile (i.e. having lower RMC) on the softer substrate (Fig. 3d). On stiffness gradients, cellular protrusions (modules) displayed higher average traction on soft than on stiff regions (Fig. 3g‒i). The cells also migrated preferentially toward the softer side, recapitulating the behavior observed in U-251MGs *in vitro* (Fig. 3j). By altering the range of the gradient, such that the side associated with higher predicted traction was the stiffer one, durotaxis could be reversed and cells clustered primarily in the stiff regions (Fig. S6a–d).

We verified the generality of these principles by applying them to model axonal pathfinding in neuronal development and regeneration (Figs. S7, S8; Supplementary Text 3). Indeed, the tendency for *Xenopus* retinal ganglion cells to grow toward softer tissue is closely analogous to negative durotaxis (*23*). Neurite elongation and pathfinding via the actin-rich neuronal growth cone (GC) at the distal end of the axon involves contractile filopodia of variable length and orientation (Fig. S7a). Applying our model to individual filopodia (Fig. S7b) and to GCs with multiple filopodia (Fig. S8a), we found that the protrusions elongated faster and generated more traction on soft substrates (0.01‒0.1 pN nm^−1^) (Fig. S7c‒h). This was consistent both with earlier predictions of relatively low optimal stiffness for neurons (*17*, *38*, *39*), and with our hypothesis that positive and negative durotaxis are governed by motor-clutch dynamics in concert with optimal stiffness. The results also suggested that gradient strength may further increase propensity for negative durotaxis: GCs steered to more compliant regions on substrates with stronger gradients (reaching a maximum at ~10 pN nm^−1^/20 μm), but did not change direction on mild gradients (~0.1 pN nm^−1^/20 μm) or on substrates that were overall stiff compared to the optimum (>1 pN nm^−1^) (Fig. S8c–e).

While U-251MGs and neurons exhibit biphasic traction forces in the physiological stiffness range, many adherent cell types do not (*11*, *18*, *40*, *41*). Rather, their traction increases as a function of substrate stiffness unless talin- and vinculin-mediated FA formation is disrupted, e.g. by depletion of both talin isoforms (Fig. 4a)(*18*). Therefore, we hypothesized that targeting adhesion reinforcement can generate an intermediate optimal stiffness and enable negative durotaxis in cell types that normally undergo only positive durotaxis. To test this, we used siRNAs to reduce talin-1 and talin-2 expression in MDA-MB-231 cells that exert increasing traction with increasing substrate stiffness (*41*) and undergo positive durotaxis in the 2–18 kPa range (*8*). Talin knockdown (Fig. 4b) resulted in significantly fewer and smaller FAs (Fig. 4c–e) and reduced traction on ~20 kPa substrates, where adhesion reinforcement is expected to counteract clutch dissociation by rapidly accumulating forces (Figs. 4f–h, S9a). EdU incorporation increased from 0.5 to 9.6 kPa and plateaued thereafter, with and without talin silencing (Fig. S9b–c). While control MDA-MB-231s seeded on 0.5–22 kPa stiffness gradients migrated toward the stiffest regions available, talin-low MDA-MB-231s phenocopied the negative durotaxis seen in U-251MGs and clustered predominantly in regions of intermediate stiffness (Fig. 4i‒j). Thus, the familiar positive durotactic behavior can be converted to negative durotaxis by manipulating the adhesive and contractile machinery of a cell to change its optimal stiffness.

**Figure 4.**
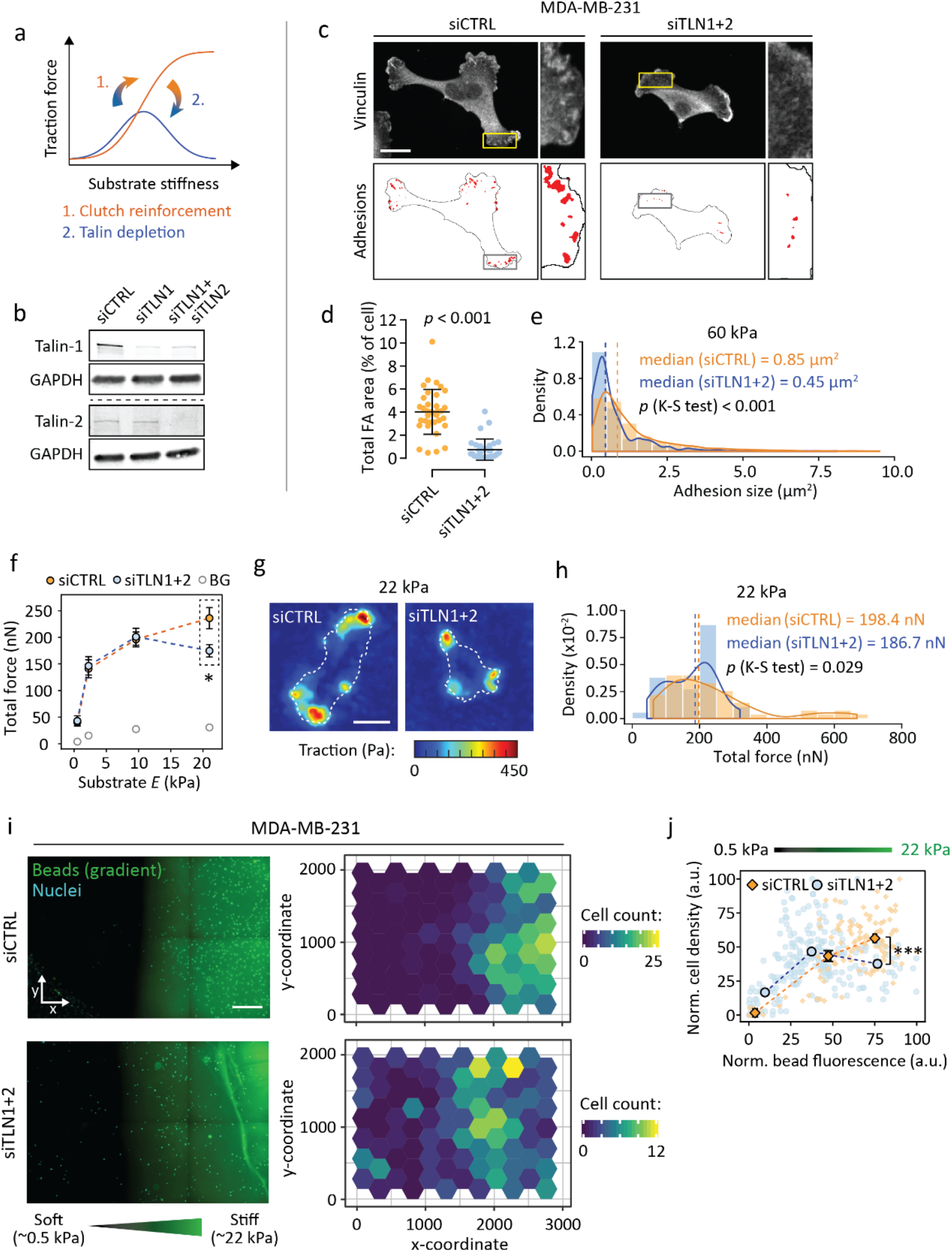
Lowering traction optimum by blocking adhesion reinforcement shifts cells from positive to negative durotaxis. (**a**) Schematic representation of the relationship between traction forces, substrate stiffness and talin/vinculin-mediated ‘clutch reinforcement’. Depletion of these clutch components forces some cell types back into a biphasic traction regime (*18*). (**b**) Representative western blot depicting talin-1 and talin-2 double knockdown in MDA-MB-231 cells. (**c–d**) Immunofluorescence images (c) and quantification (d) of focal adhesions in MDA-MB-231s on 60 kPa substrate, without and after talin knockdown. Scale bar, 20 μm. Mean ± SD of n = 32‒35 cells, analyzed by Mann-Whitney test. (**e**) Distribution of focal adhesion sizes in control and talin-low cells. Histograms overlaid with probability density functions, dashed lines indicate medians. n = 524‒1844 adhesions from 32‒35 cells, analyzed by Kolmogorov-Smirnov test. Representative of two independent experiments. (**f–h**) Traction force analysis of control and talin-low MDA-MB-231s. (f) Total force exerted by the cells as a function of substrate stiffness. Background, BG. Mean ± SEM of n = 18‒55 cells from three independent experiments, *p = 0.029. (g) Representative traction maps from cells on 22 kPa substrate. Cell outlines are indicated by white dashed lines. Scale bar, 20 μm. (h) Histograms of the 22 kPa data overlaid with probability density functions, dashed lines indicate medians. n = 37‒55 cells from three independent experiments, analyzed by Kolmogorov-Smirnov test. (**i**) (Left) Representative regions of two 0.5‒ 22 kPa polyacrylamide stiffness gradients, 72 hours after being seeded with MDA-MB-231 cells (indicated by nuclear staining). Scale bar, 500 μm. (Right) Quantification of cells across the gradients. (**j**) Relative cell densities in different parts of the gradients, overlaid with binned data. Mean ± SEM of n = 13‒141 ROIs per bin, from one (siCTRL) or two (siTLN1+2) gradient gels, representative of three independent experiments. Analyzed by Mann-Whitney test.

The concept of cells moving toward environments where they can exert more traction is intuitive, but has been previously understood in the context of denser, stiffer ECM providing cells with more stable anchorage (*7*). Our results demonstrate the additional capacity of individual cells to migrate toward softer environments, i.e. negative durotaxis, which can be explained by a motor-clutch-based model. Cells that lack robust adhesion reinforcement, such as U-251MG glioma cells or talin-low MDA-MB-231 breast cancer cells, tend to exert maximal traction on substrates of intermediate stiffness, and migrate along gradients to reach this optimum by positive or negative durotaxis (Fig. S10). The same mechanism is likely to contribute to the recently described neurite growth toward soft matrix (*23*).

Besides directly reinforcing connections to stiff matrix, mechanosensitive FA formation may promote positive durotaxis by additional mechanisms. Preferential trafficking of adhesion components toward existing FAs (*42*), local activation of mechanically gated ion channels (*43*) or other biochemical signaling pathways initiated at the FAs (*34*) may all contribute to further polarization of cell-matrix adhesion and, consequently, of cellular traction forces. How these factors influence stiffness optima on different substrates, and in different biological conditions, will be an interesting topic for future research. Taken together, our results point to a single, conserved mechanism for stiffness sensing and durotaxis across a broad range of cell types, with motor-clutch dynamics driving traction generation and choices between positive and negative durotaxis.

---

**Figure S1.**
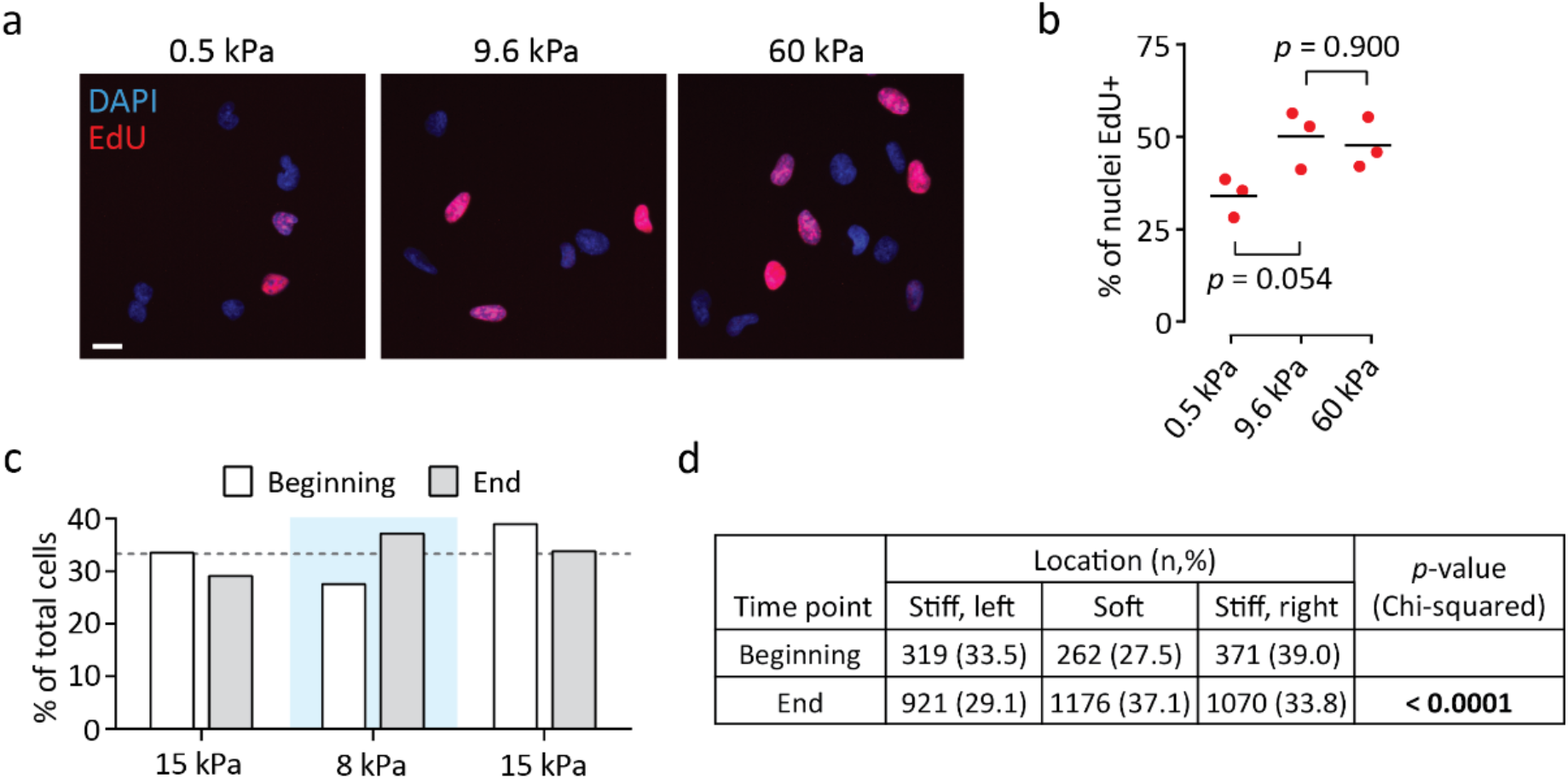
Mechanosensitivity of U-251MG proliferation and clustering on stepwise stiffness gradients. (**a**–**b**) Fluorescence images (a) and quantification (b) depicting EdU incorporation by U-251MG cells on 0.5*‒*60 kPa substrates. Scale bar, 20 μm. Mean values from three independent experiments. Analyzed by one-way ANOVA and Sidak’s *post hoc* test. (**c**–**d**) Total number of cells in the different gradient regions in (Fig. 1e and g). (c) Bar graph, n = 952‒3,167 cells per time point. (d) Contingency table summarizing the data, analyzed by chi-squared test.

**Figure S2.**
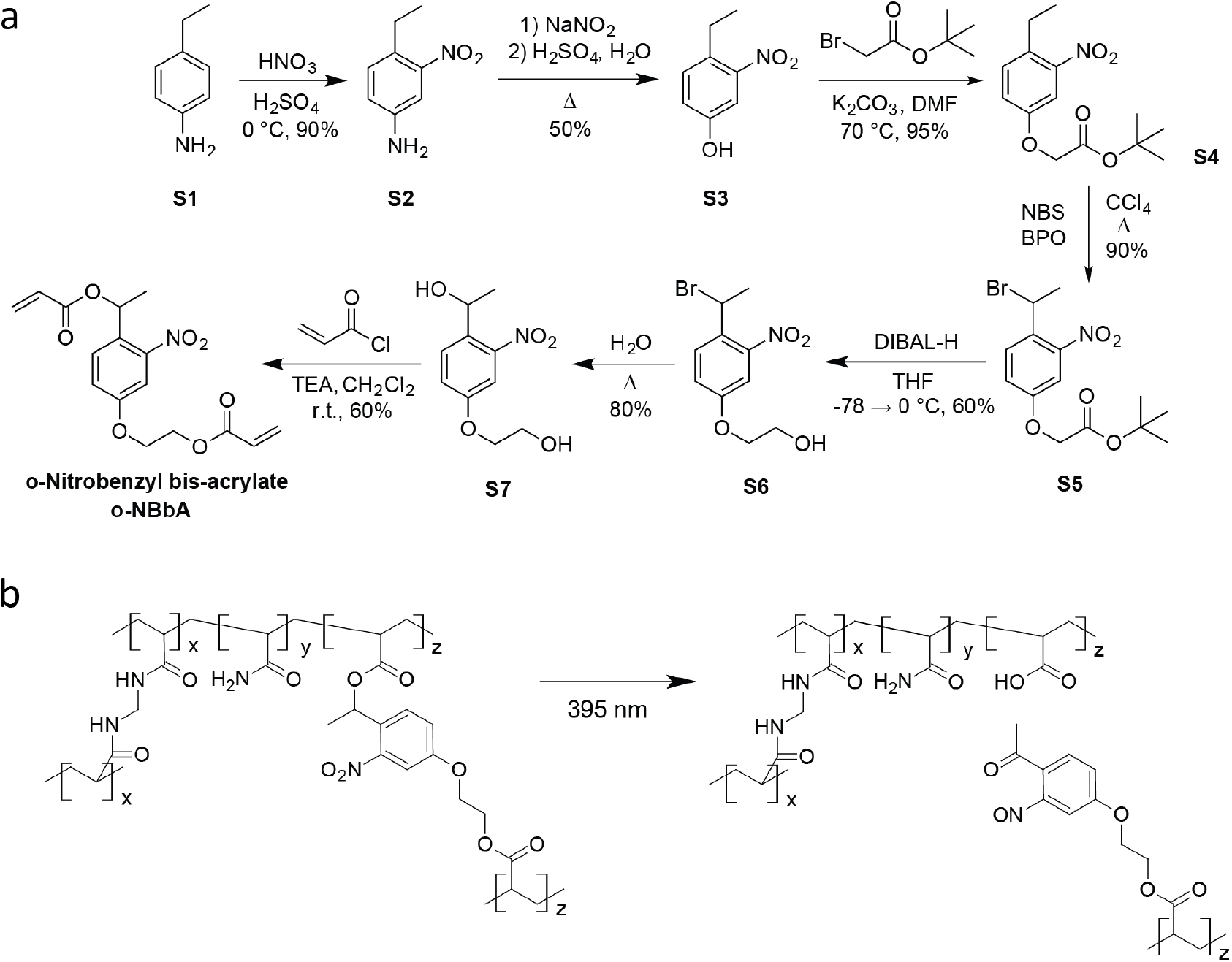
Synthesis and photochemistry of o-nitrobenzyl bis-acrylate. (**a**) Schematic of the synthesis of o-NBbA. (**b**) Copolymerization of o-NBbA with acrylamide and bis-acrylamide yields hydrogels composed of strands of polyacrylamide crosslinked by either o-NBbA or bis-acrylamide. UV irradiation cleaves the photolabile o-NBbA, resulting in gels with lower crosslinking density and hence lower stiffness. The process does not release any byproducts to the gel environment.

**Figure S3.**
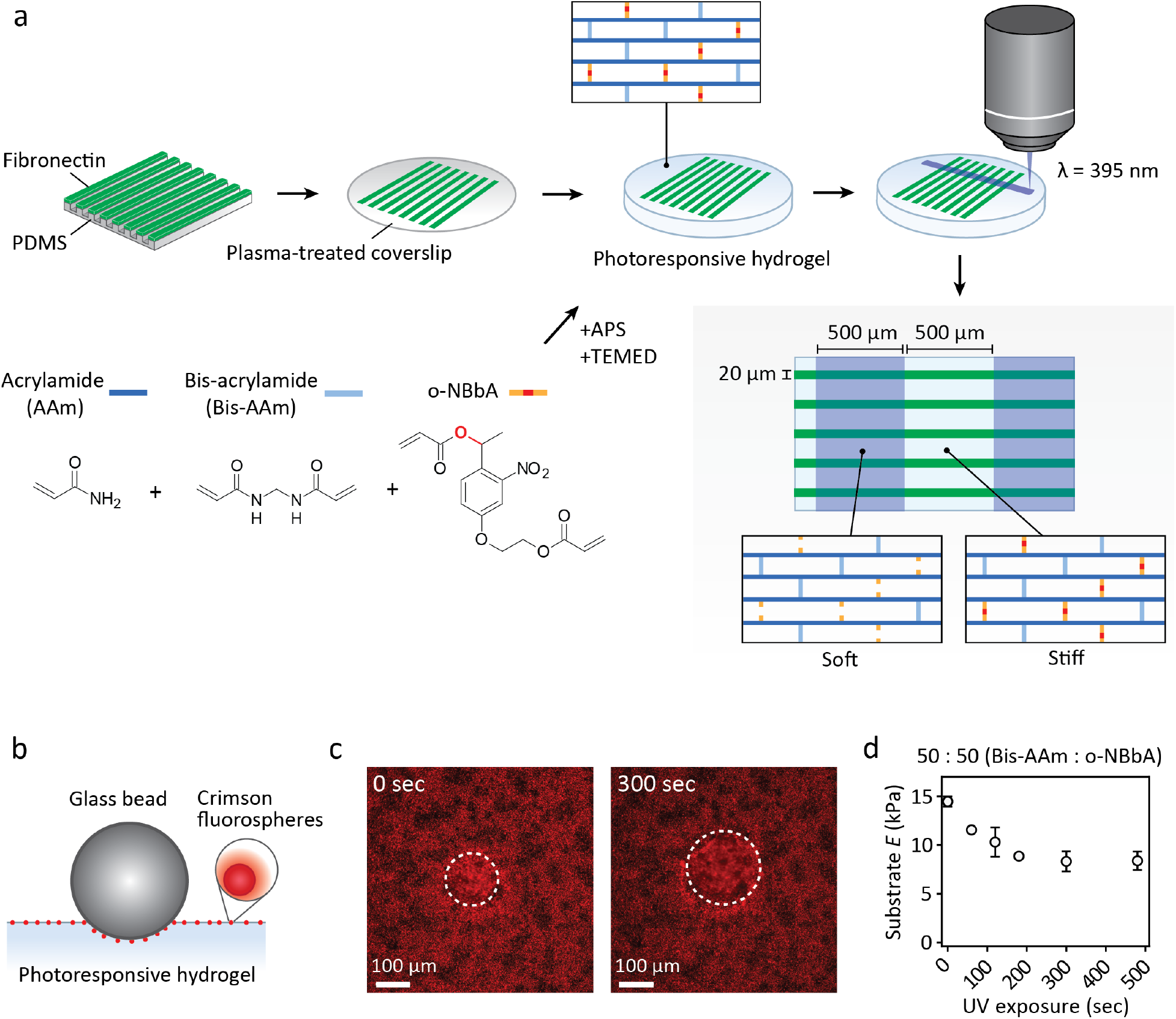
Preparation and characterization of photoresponsive hydrogels. (**a**) Schematic representation of stiffness gradient preparation for migration experiments. The photocleavable carbon-oxygen bond in o-NBbA is indicated by red color. (**b–d**) Stiffness characterization by bead indentation. A schematic representation of the technique (b), representative fluorescence images (c) and quantified results (d) from the experiment, depicting hydrogel elasticity as a function of UV exposure. Dashed lines indicate indented, out-of-focus areas in the gel. Mean ± SD of n = 3 measurements.

**Figure S4.**
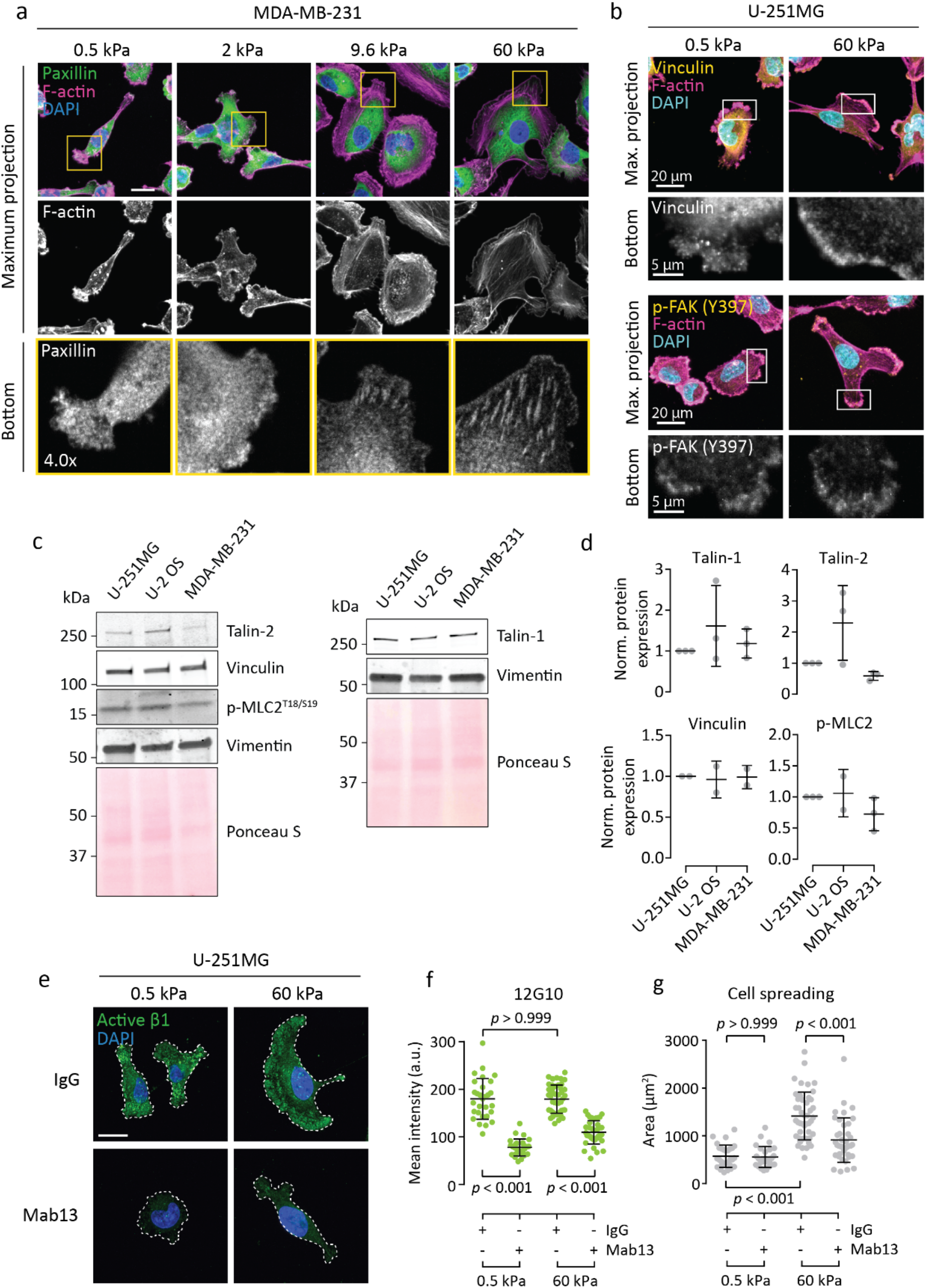
Focal adhesion maturation and adhesion components in U-251MG and other cancer cells. (**a**) Immunofluorescence images of paxillin and F-actin in MDA-MB-231 cells on 0.5‒60 kPa substrates. The bottom panels show individual focal planes from confocal stacks, corresponding to the basal side of each cell. Scale bar, 20 μm. (**b**) Immunofluorescence images of vinculin (top), p-FAK (bottom) and F-actin in U-251MG cells on 0.5 and 60 kPa substrates. The bottom panels show individual focal planes from confocal stacks, corresponding to the basal side of each cell. (**c–d**) Representative western blots (c) and quantification (d) depicting talin-1/2, vinculin and p-MLC2 levels across three different cell lines. Densitometric measurements were normalized to vimentin, mean ± SD of 2‒3 independent experiments. (**e–f**) Immunofluorescence images (e) and quantification (f) showing active β1-integrin (clone 12G10) in U-251MGs on 0.5 and 60 kPa substrates. The cells were treated with a control antibody (normal rat IgG) or β1 function-blocking Mab13 for two hours before fixation. Mean ± SD of n = 27–45 cells, analyzed by Kruskal-Wallis one-way ANOVA and Dunn’s *post hoc* test. Representative of two independent experiments. (**g**) Spreading of U-251MGs on 0.5 and 60 kPa substrates, without or after β1-integrin blocking by Mab13. Mean ± SD of n = 27‒45 cells, analyzed by Kruskal-Wallis one-way ANOVA and Dunn’s *post hoc* test. Representative of two independent experiments.

**Figure S5.**
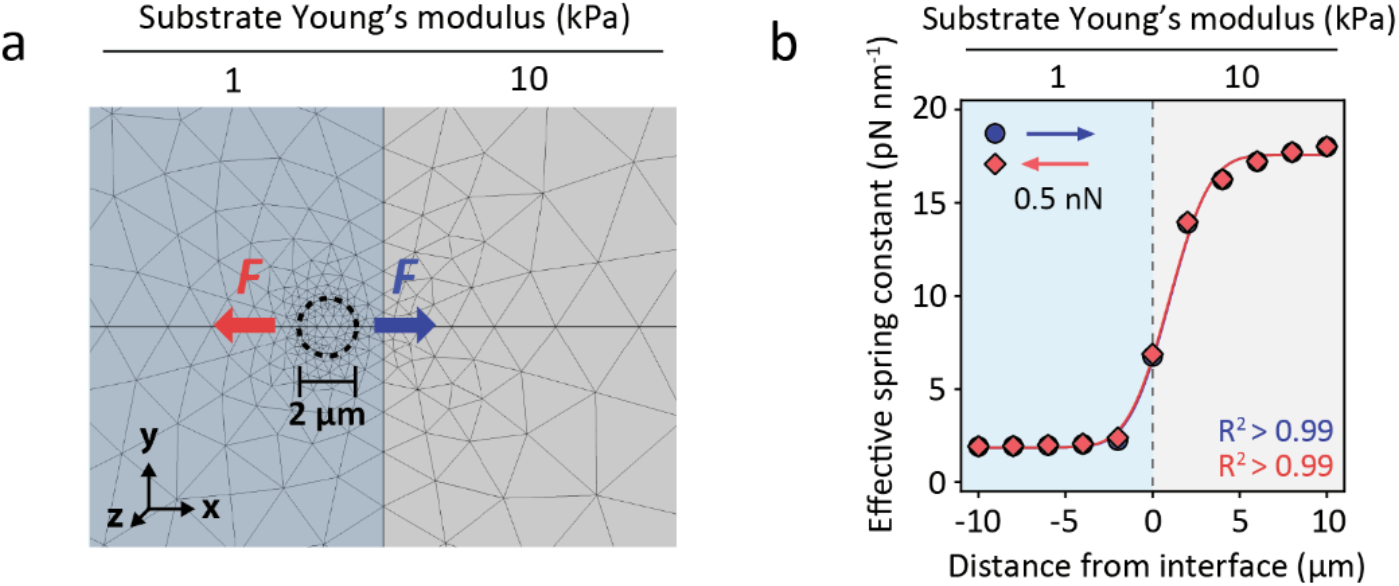
Finite element analysis of polyacrylamide displacement next to a stepwise elastic gradient. (**a**) COMSOL Multiphysics® model setup (**b**) The effect of steep elastic gradients on the effective spring constant of polyacrylamide. A lateral 0.5 nN force was exerted on the substrate through a circular adhesion zone (*r* = 1 μm) as shown in (a). The position of the adhesion zone was adjusted repeatedly at 2 μm steps. The direction of the force was varied by 180° but was always parallel to the gradient. In both cases, normal cumulative distribution function was a good fit to the data.

**Figure S6.**
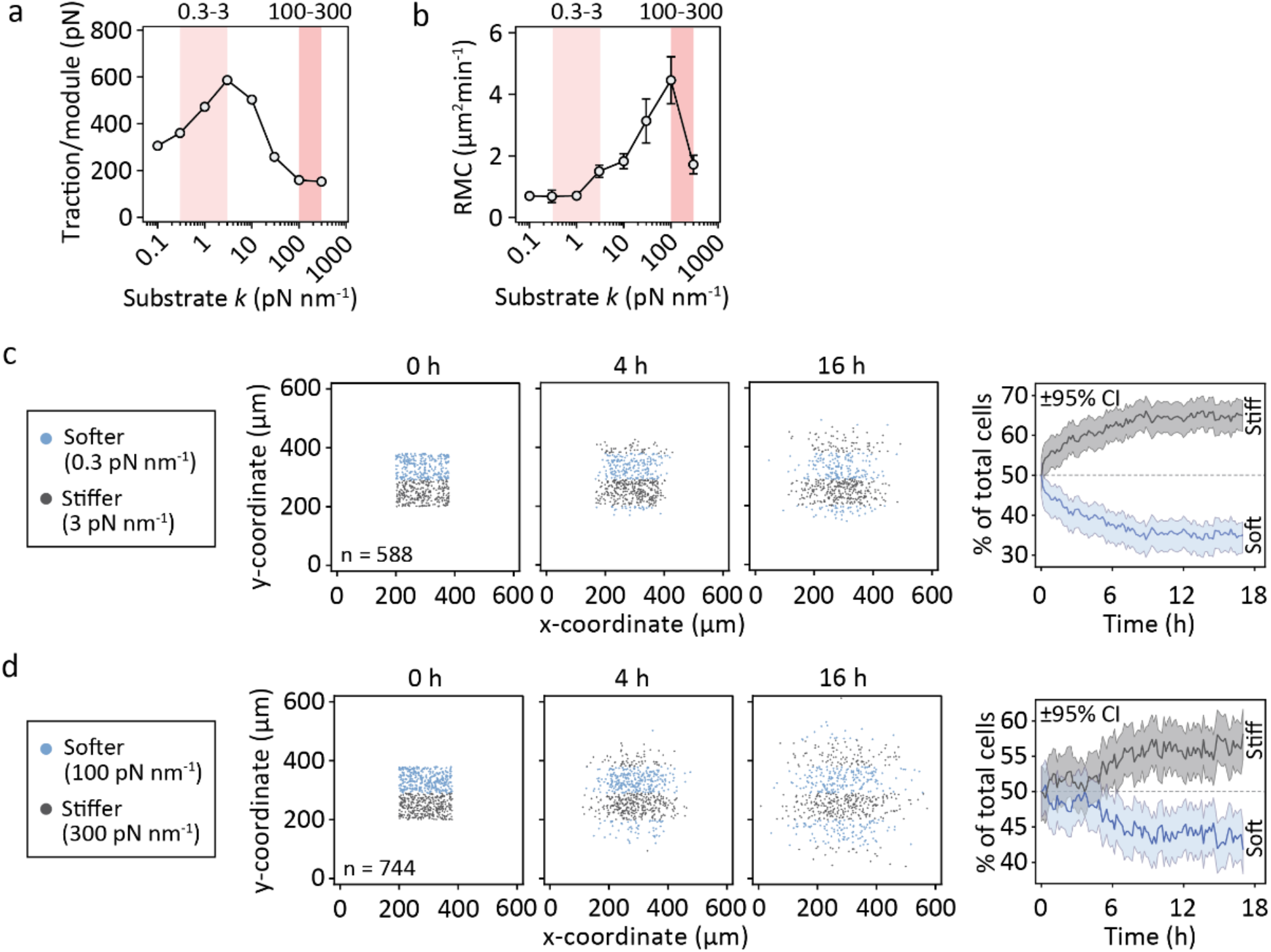
Modifying the range of the gradient can reverse durotaxis *in silico*. (**a–b**) Module traction forces (a) and RMC (b) of the simulated cells as a function of substrate stiffness, as in (Fig. 3c-d). Red overlays highlight the ranges of the 0.3‒3 pN nm^−1^ and 100‒300 pN nm^−1^ gradients in (c‒d). Mean ± SEM of n = 10 cells. (**c–d**) Evolution of cell density on mechanically heterogeneous substrates over time. (c) Coordinates of individual cells on the 0.3‒3 pN nm^−1^ gradient 0, 4 and 16 hours into the simulation (left) and the fraction of cells residing in the stiffer and softer areas over the course of the simulation (right). ±95% CI, n = 588 cells. (d) As above, but for the 100‒300 pN nm^−1^ gradient. n = 744 cells.

**Figure S7.**
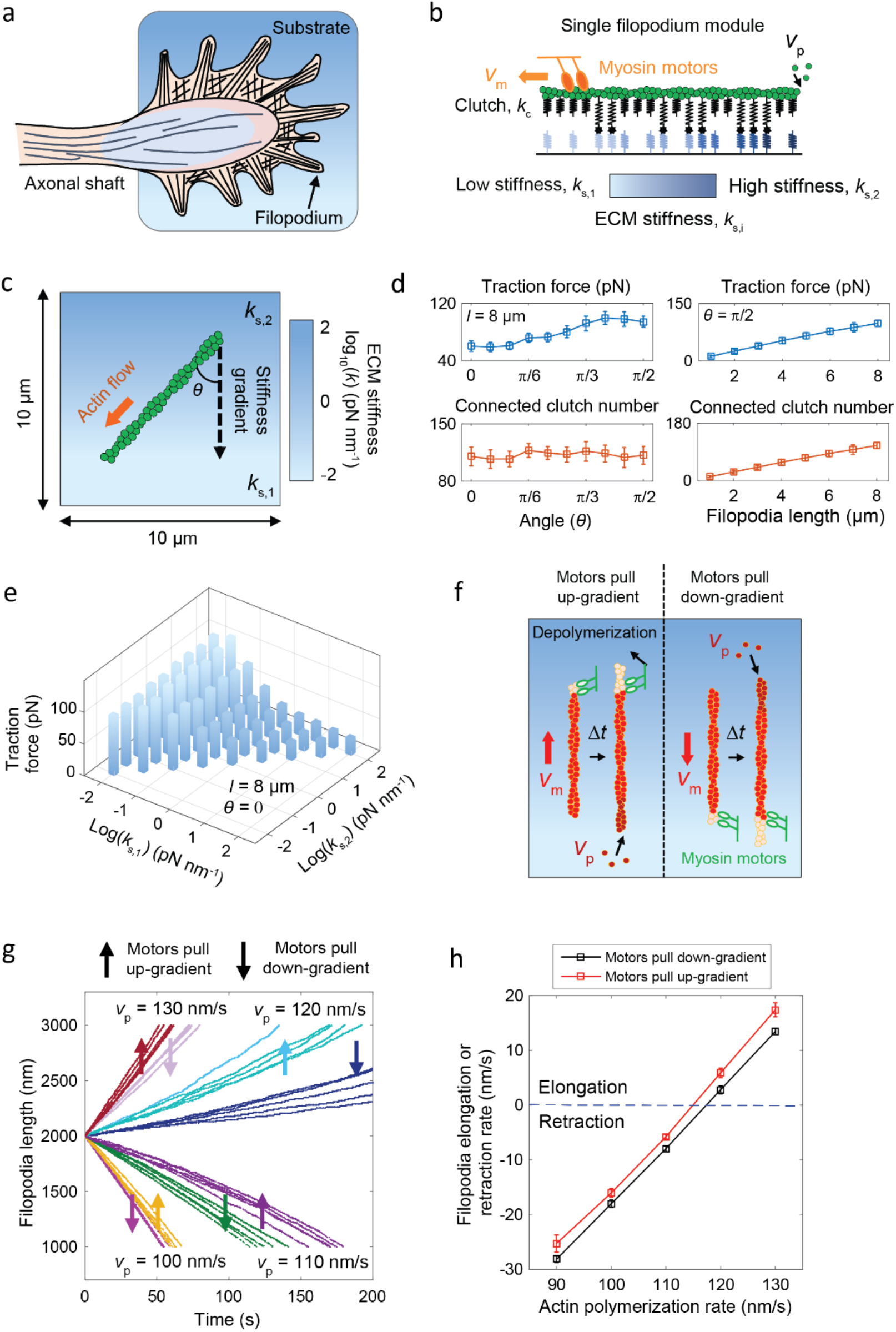
Motor-clutch model of filopodial dynamics. (**a**) Schematic representation of a neuronal GC. Filopodia, surrounded by a less polarized actin network, reside in the peripheral domain. They are separated from the axon by a thin transitional domain, and a central domain (light blue) that is primarily composed of microtubules. (**b**) The filopodia in GCs are modeled as individual motor-clutch modules, with adhesion springs (homogeneous stiffness), substrate springs (heterogeneous stiffness) and inward actin flow resulting from active myosin motors. Actin monomers are added into the filaments at a constant rate. (**c**) Setup used in the single-filopodium simulations. The filopodium interacts with the substrate in a set orientation relative to the linear stiffness gradient. (**d**) (Left) Traction force exerted by the filopodium increases when the protrusion is pointing down the gradient, toward softer substrate. (Right) Perpendicular to the gradient, traction increases with filopodia length mainly due to more clutches being available to bind with the substrate. Data shown are from *n* = 10 independent simulations. (**e**) Average traction exerted by a single filopodium on different substrate stiffness gradients. Data represent means of *n* = 10 simulations. (**f**) Filopodia length is affected by both actin flow, *v*_*m*_, and the polymerization rate, *v*_*p*_. Depending on the orientation of the filopodium, the actin may flow toward soft (filopodium pointing up the gradient) or stiff (filopodium pointing down the gradient) substrate. (**g**) Evolution of filopodia length on stiffness gradients upon different actin polymerization rates. The different combinations of *v*_*p*_ and filopodia orientation are color-coded, while each line represents the temporal variation in the length of a single filopodium. (**h**) Effect of actin polymerization and orientation relative to a stiffness gradient on the filopodia elongation/retraction rate. Mean ± SEM in (d) and (h).

**Figure S8.**
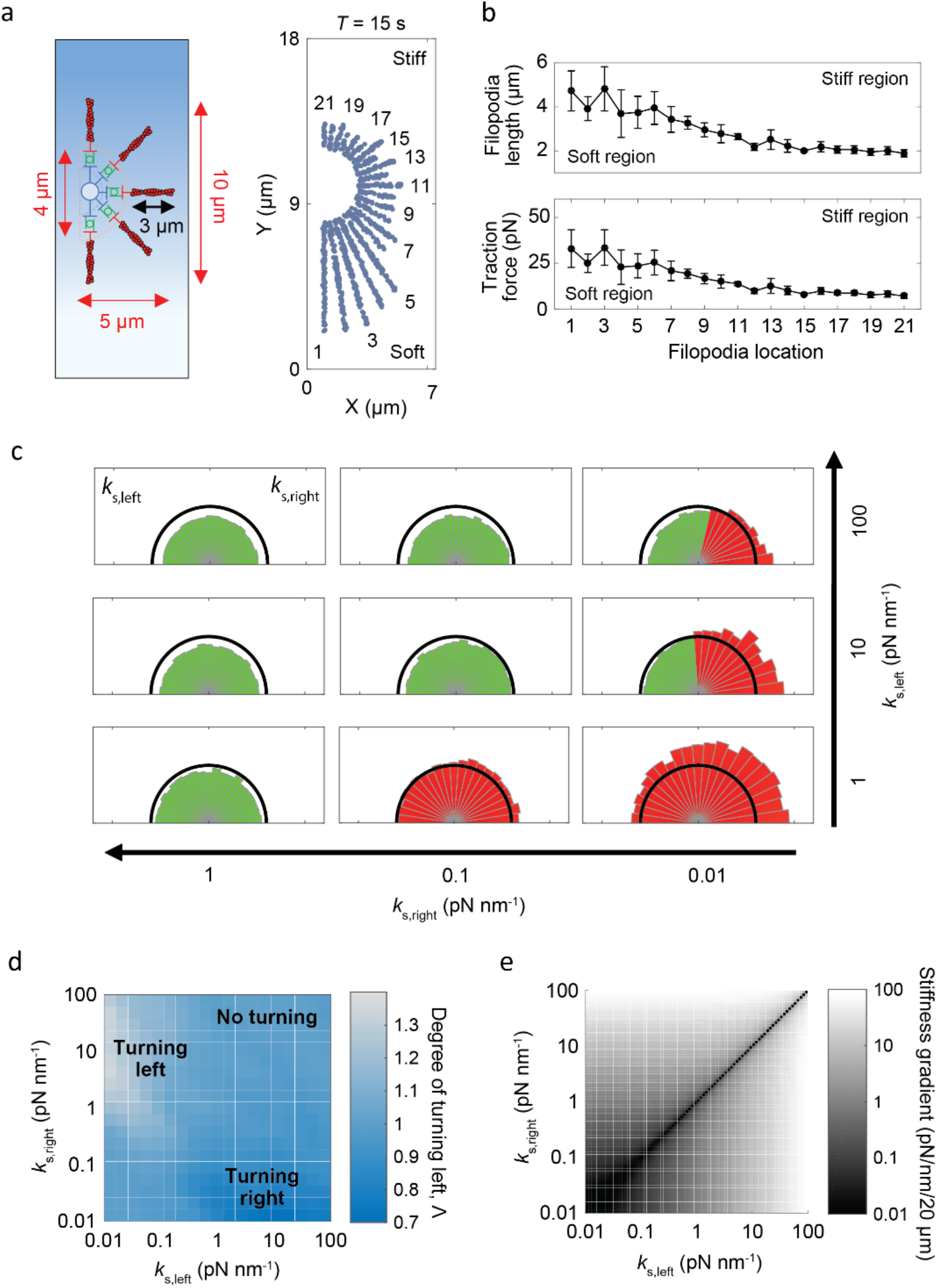
Motor-clutch model predicts growth cone steering toward soft matrix. (**a**) Schematic representation of the GC model. (Left) Dimensions of a newly initialized GC. (Right) Each GC consists of multiple filopodia, distributed between –π/2 and π/2 relative to the axon. On stiffness gradients (*K*_*s*,1_ = 0.01 pN nm^−1^, *K*_*s*,2_ = 100 pN nm^−1^), filopodia on the more compliant side of the substrate rapidly outgrow the others, leading to effective turning of the GC. (**b**) Filopodia length (top) and traction (bottom) based on their orientation around the GC central domain. On stiffness gradients, filopodia pointing toward the softer substrate elongate faster and generate more traction. Data shown are from n = 10 independent simulations. (**c**) Examples of GC behavior on different stiffness gradients. Green denotes filopodia that are retracting during the course of the simulation, red denotes filopodia that are elongating. Depending on the gradient, individual GCs may retract or enlarge isotropically, or steer toward the softer substrate. Displayed are means of n = 10 simulations. (**d**) Phase diagram of GC turning to left, Δ, on different mechanically graded substrates. (**e**) Phase diagram depicting the strength of the stiffness gradient for varying *K*_*s*,1_ and *K*_*s*,2_. Gradient strength alone cannot explain the magnitude of Δ, if the whole substrate is stiffer than the optimal range for individual filopodia (Fig. S7e).

**Figure S9.**
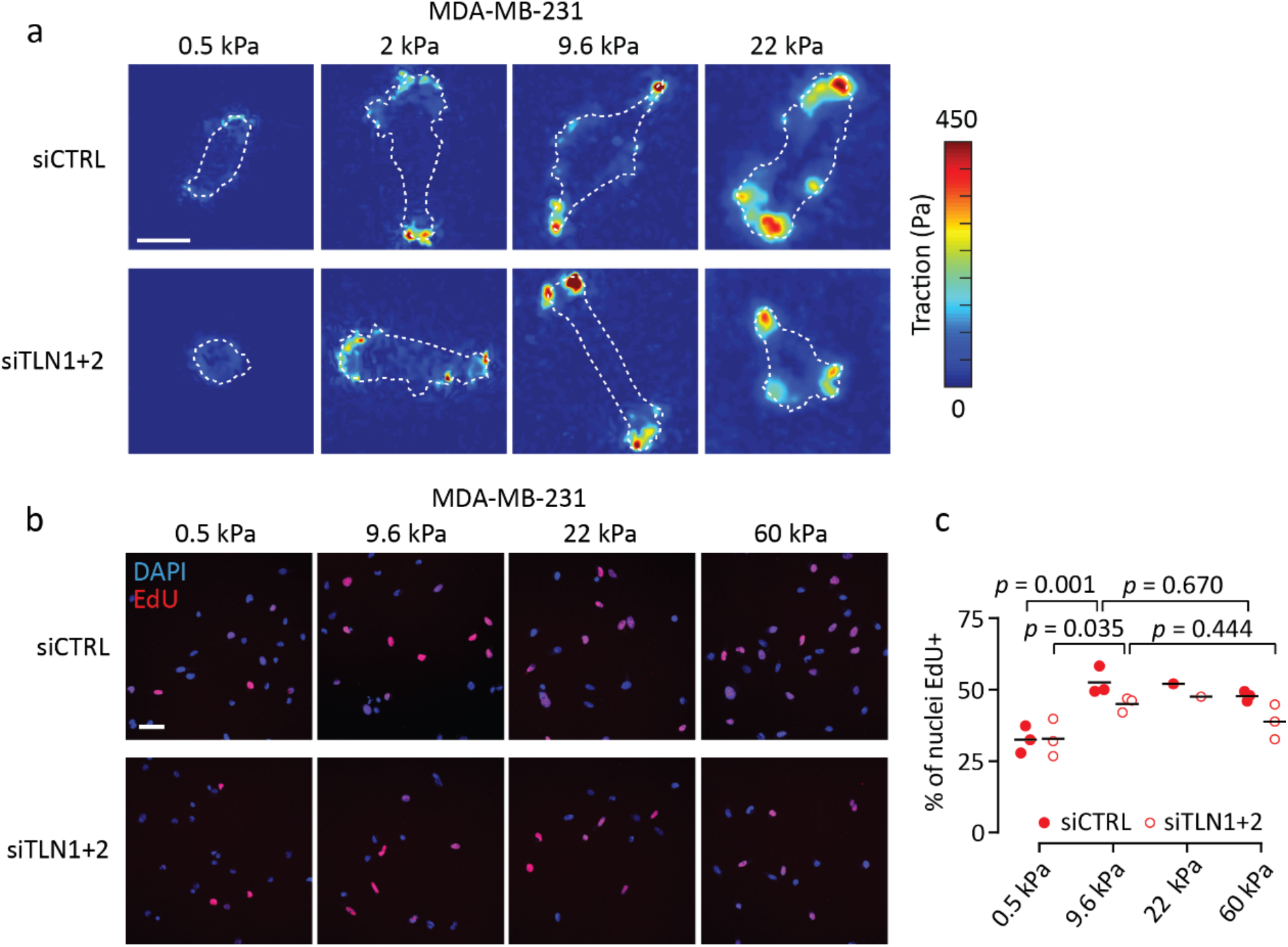
Mechanosensitive traction and proliferation of MDA-MB-231 cells. (**a**) Representative traction maps from MDA-MB-231 cells on 0.5*–*22 kPa substrates, corresponding to the data in (Fig. 4f). Cell outlines are indicated by white dashed lines. Scale bar, 20 μm. (**b***–***c**) Fluorescence images (a) and quantification (b) depicting EdU incorporation by control and talin-low MDA-MB-231 cells on 0.5*–*60 kPa substrates. Scale bar, 50 μm. Mean values from one to three independent experiments. Analyzed by one-way ANOVA and Sidak’s *post hoc* test.

**Figure S10.**
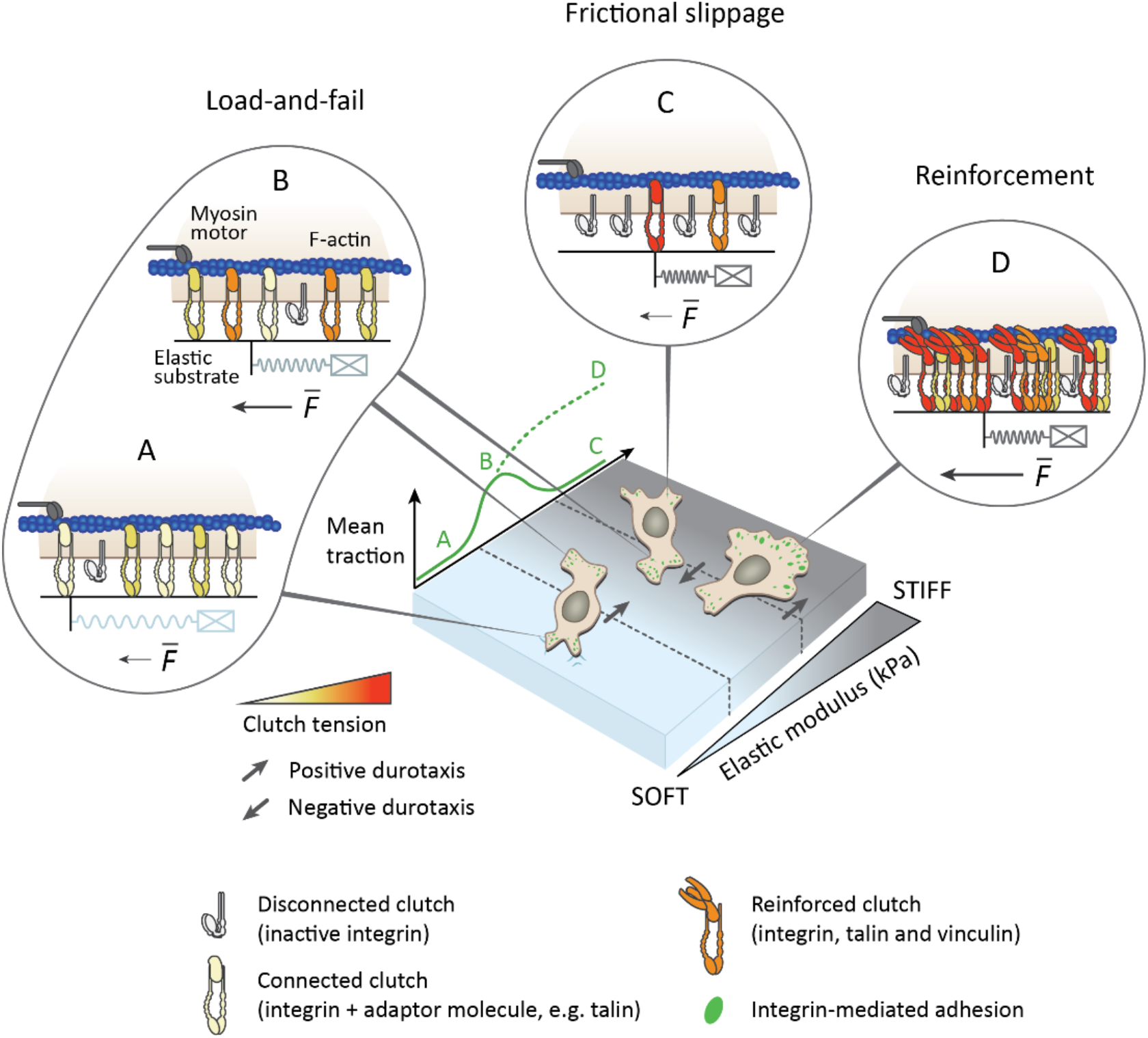
Schematic representation of the regulation of positive and negative durotaxis by motor-clutch dynamics. Cell-intrinsic molecular machinery dictates the cell’s capacity to exert force on mechanically heterogeneous substrates, driving positive or negative durotaxis. Without clutch reinforcement (mechanosensitive FA formation, D), the motor-clutch model predicts a biphasic relationship between traction force and substrate stiffness (A*–*C) (*17*, *18*, *25*–*29*). This fundamental relationship, and the physical reinforcement of cell-matrix adhesion by FAs, are likely to be further influenced by biochemical signaling pathways and feedback loops that modulate the expression, activity and localization of individual cytoskeletal and clutch components, in a cell type-dependent manner.

## Materials and Methods

### Cell culture and transfections

U-251MG human glioblastoma cells were obtained from Dr. G. Yancey Gillespie (U. Alabama-Birmingham), authenticated using a short tandem repeat assay (University of Arizona Genetics Core) and cultured in Dulbecco’s modified Eagle’s medium (DMEM)/F-12 (Gibco, 11320-074) supplemented with 8% fetal bovine serum (Sigma, F7524). MDA-MB-231 human breast adenocarcinoma cells were purchased from American Type Culture Collection and authenticated using a short tandem repeat assay (Leibniz Institute DSMZ ‒ German Collection of Microorganisms and Cell Cultures, Braunschweig, Germany). U-2 OS human osteosarcoma cells were acquired from DSMZ. Both MDA-MB-231 and U-2 OS were cultured in high-glucose DMEM (Sigma, D5796-500ML) supplemented with 10% fetal bovine serum (Sigma, F7524), 2 mM L-glutamine (Sigma, G7513-100ML) and 1x non-essential amino acids (Sigma, M7145-100ML). The cells were tested for mycoplasma contamination and cultured at +37 °C, 5% CO_2_ in a humidified incubator.

For transient downregulation of target proteins, the cells were transfected with corresponding siRNAs at a 50 nM concentration per oligo. The transfections were conducted using Opti-MEM Reduced Serum Medium (Thermo Fisher Scientific, 31985-047) and Lipofectamine RNAiMAX reagent (Thermo Fisher Scientific, 56532) according to the manufacturer’s instructions. The siRNAs used were Hs_TLN1_3 FlexiTube siRNA (Qiagen, SI00086975), Hs_TLN2_3 FlexiTube siRNA (Qiagen, SI00109277) and AllStars Negative Control siRNA (Qiagen, 1027281). Silenced cells were grown for 24 (beginning of migration experiments) to 72 hours before they were used for experiments.

### Antibodies

The following antibodies were used at the indicated dilutions: ms anti-paxillin (BD Biosciences, 612405, 1:200 for IF), rbt anti-paxillin (Santa Cruz Biotechnology, sc-5574, 1:200 for IF), ms anti-vinculin (Sigma, V9131, 1:200 for IF, 1:1000 for WB), ms anti-talin-1 (Novus, NBP2-50320, 1:1000 for WB), ms anti-talin-2 (Novus, NBP2-50322, 1:1000 for WB), ms anti-FAK (BD Biosciences, 610088, 1:1000 for WB), rbt anti-p-FAK (Y397) (Cell Signaling Technology, 8556, 1:100 for IF, 1:1000 for WB), rbt anti-MLC2 (Cell Signaling Technology, 3672, 1:1000 for WB), rbt anti-p-MLC2 (T18/S19) (Cell Signaling Technology, 3674, 1:1000 for WB), rbt anti-ERK1/2 (Cell Signaling Technology, 9102, 1:1000 for WB), rbt anti-p-ERK1/2 (T202/Y204) (Cell Signaling Technology, 4370, 1:1000 for WB), ms anti-YAP (Santa Cruz Biotechnology, sc-101199, 1:200 for IF), rbt anti-vimentin (Cell Signaling Technology, 5741, 1:1000 for WB), ms anti-GAPDH (HyTest, MAb 6C5, 1:5000 for WB), ms anti-active β1-integrin (clone 12G10, in-house production, 5 μg/ml for IF), rat anti-inactive β1-integrin (clone Mab13, in-house production, 10 μg/ml for cell culture), and normal rat IgG2a kappa isotype control (eBioscience, 14-4321-85, 10 μg/ml for cell culture).

Additionally, the following secondary antibodies were used for immunofluorescence and immunoblots at the indicated dilutions: Alexa Fluor 488/568-conjugated secondary antibodies raised against mouse (Invitrogen, A21202 and A10037, 1:400 for IF) and rabbit (Invitrogen, A21206 and A10042, 1:400 for IF), IRDye 800CW Donkey anti-Mouse IgG (LI-COR Biosciences, 926-32212, 1:5000 for WB), IRDye 800CW Donkey anti-Rabbit IgG (LI-COR Biosciences, 926-32213, 1:5000 for WB), and IRDye 680LT Donkey anti-Mouse IgG (LI-COR Biosciences, 926-68022, 1:5000 for WB).

### EdU incorporation assay

To measure the rate of EdU (5-ethynyl-2′-deoxyuridine) incorporation into DNA, cells were grown on hydrogels for 24 hours, after which they were prepared into fluorescence microscopy samples using an EdU Proliferation Assay Kit (Abcam, ab222421), according to the manufacturer’s instructions. Briefly, the cells were supplemented with 20 μM EdU for 2 hours, fixed, permeabilized, and the EdU was stained with iFluor 647 azide via a copper-catalyzed click reaction. Nuclei were counterstained before imaging (see below).

### Blocking β1-integrin function with antibodies

U-251MG cells were grown on 0.5 kPa and 60 kPa hydrogels for 24 hours, after which they were treated with 10 μg/ml of anti-inactive β1-integrin (i.e. function-blocking) clone Mab13 or normal rat isotype control for 2 hours (see the list of antibodies for details). The cells were fixed and processed for immunofluorescence imaging.

### Cell migration on stiffness gradient substrates

For analysis of cell migration on continuous 2D stiffness gradients, 15,000 (MDA-MB-231)‒20,000 (U-251MG) cells were seeded on a fibronectin-functionalized stiffness gradient hydrogel. Even distribution of cells in the beginning of the experiment was confirmed visually (via brightfield microscopy) and by recording the positions of individual nuclei along the gradient using SiR-DNA. The plate was returned to the incubator for 48 (U-251MG) or 72 hours (MDA-MB-231), after which the cells were fixed and nuclei were re-visualized with DAPI.

For live imaging of U-251MG migration on stepwise gradient hydrogels, 10,000 cells were seeded per dish and allowed to settle in the incubator for 30 min prior to imaging. Time-lapse movies were acquired at 20 or 30 min intervals for 45 to 60 hours. The number of cells in the soft and stiff regions of the gel, in the beginning and end of the experiment, was quantified. Additionally, the movies were analyzed for cells directly on top of a stiffness gradient. Such cells were tracked over time to investigate their bias for migrating toward either stiffness. Mitotic, dying or crowded cells were excluded from the analysis.

### Western blotting

Cells on hydrogels were placed on ice, rinsed twice with ice-cold PBS and scraped into lysis buffer [50 mM Tris-HCl pH 7.5, 150 mM NaCl, 1% SDS, 0.5% Triton X-100, 5% glycerol, supplemented with protease (Roche, 05056489001) and phosphatase (Roche, 04906837001) inhibitors]. The lysates were vortexed, placed on a heat block (+90 °C) for 10 min and sonicated before separation by SDS-PAGE (4-20% Mini-PROTEAN TGX Gels, Bio-Rad, 456-1096). Next, the proteins were transferred to nitrocellulose membranes and visualized using 1% Ponceau S staining solution. The membranes were blocked with 5% skimmed milk in TBST and incubated with the indicated primary antibodies overnight at +4°C, followed by fluorophore-conjugated secondary antibodies for 1 to 2 hours at room temperature. All the antibodies were diluted in StartingBlock blocking buffer (Thermo Fisher Scientific, 37538). Finally, the membranes were scanned using an Odyssey infrared imaging system (LI-COR Biosciences).

### Conventional polyacrylamide hydrogels

Glass-bottom dishes (Cellvis, D35-14-1-N) were treated for 20 min at room temperature with 100 μl of Bind-Silane solution – a mixture of 3-(trimethoxysilyl)propylmethacrylate (7.15% by volume, Sigma-Aldrich, M6514) and acetic acid (7.15% by volume) in absolute ethanol – to promote gel attachment to the glass surface. After the Bind-Silane was aspirated, the glass was washed twice with ethanol and left to dry completely. For homogeneous (constant Young’s modulus) hydrogels, pre-defined ratios of 40% (w/v) acrylamide (Sigma-Aldrich, A4058) and 2% (w/v) N,N-methyl-bis-acrylamide (Sigma-Aldrich, M1533) were mixed in PBS on ice and vortexed carefully. The final concentrations were adjusted to yield a desired Young’s modulus (Table S1). Gels that were indicated for traction force microscopy were supplemented with additional 0.2 μm yellow-green fluorescent (505/515) microspheres (~1.5 × 10^10^/ml final concentration, Invitrogen, F8811), which were sonicated for 3 min prior to use. Polymerization was initiated by addition of 10% ammonium persulfate (APS, final 0.1% by volume, Bio-Rad) and N,N,N′,N′-tetramethylethylenediamine (TEMED, final 0.2% by volume, Sigma-Aldrich, T-9281) to the solution. Immediately afterwards, 13 μl of the solution was pipetted onto the glass-bottom dish and a 13 mm circular coverslip was placed on top of the droplet. After polymerization for ~1 hour at room temperature, the gel was immersed in PBS for 5 min, the top coverslip was gently removed, and the gel was washed twice with PBS to remove any excess acrylamide. Hydrogels with continuous 2D stiffness gradients were fabricated as described previously (*30*). Briefly, 0.5 kPa and 22 kPa acrylamide prepolymer solutions were prepared and 0.1 μm fluorescent (505/515) microspheres (~1.2 × 10^11^/ml final concentration, Invitrogen, F8803) were added to the 22 kPa solution. After polymerization was initiated, the two solutions were allowed to diffuse together on a glass-bottom dish, under a glass coverslip, to yield a gradient wherein microsphere density correlates linearly with the Young’s modulus of the substrate.

Prior to use, the hydrogels were activated by a combination of 0.2 mg/ml Sulfo-SANPAH (Thermo Fisher Scientific, 22589) and 2 mg/ml N-(3-dimethylaminopropyl)-N′-ethylcarbodiimidehydrochloride (EDC, Sigma-Aldrich, 03450) in 50 mM 4-(2-hydroxyethyl)piperazine-1-ethanesulfonic acid (HEPES). 500 μl of the solution was added on top of the hydrogel and incubated for 30 min at room temperature, protected from light and subjected to gentle agitation. The gel and solution were then UV-irradiated for 10 min (28-32 mW/cm^2^) to activate the Sulfo-SANPAH, and the plate was washed with PBS three times to remove any residual compounds. Finally, each hydrogel was functionalized by incubation in 10 μg/ml fibronectin solution overnight at +4°C.

Cells that were collected for protein lysates were cultured on commercial hydrogel-coated 6-well plates (Matrigen, SW6-EC-0.5/SW6-EC-8/SW6-EC-50). These gels were similarly coated with 10 μg/ml of fibronectin before use.

### Synthesis of o-NBMA

#### 2-nitro-4-ethyl aniline (S2)

p-Ethyl aniline (5 g, 41.3 mmol) was added dropwise to a cold solution of concentrated H_2_SO_4_ (30 ml) and stirred for 5 min. In a separate flask, 5.3 ml of 70% HNO_3_ (82.6 mmol) was mixed with an equal volume of H_2_SO_4_, and added dropwise to the reaction vessel, followed by 15 min stirring at 0 °C. Thin layer chromatography (TLC) analysis (Hex:EtOAc, 2:1, v/v) indicated complete conversion to the product. The reaction was quenched by pouring the mixture into 200 ml ice water. The resulting precipitate was filtered and washed with H_2_O to yield compound **S2**(6.2 g, 90%).

^1^H NMR (500 MHz, CDCl_3_) δ ppm 1.099 (t, *J* = 7.5 Hz, 3H), 2.612 (q, *J* = 7.0 Hz, 2H), 5.558 (s, 2H), 6.804 (dd, *J* = 8.0, 2.5 Hz, 1H), 7.041 (d, *J* = 2.5 Hz, 1H), 7.095 (d, *J* = 8.5 Hz, 1 H)

^13^C NMR (400 MHz, DMSO-d6) 149.3411, 147.8646, 131.6051, 124.0586, 118.8417, 107.9194, 24.4890, 15.3093

HRMS *(m/z):* [M]^+^ calcd for [C_8_H_10_N_2_O_2_]^+^ 166.0737, found 166.0737.

#### 4-ethyl-3-nitrophenol (S3)

Compound **S2** (6.2 g, 37.3 mmol) was suspended in a mixture of H_2_SO_4_ and H_2_O (1:3, v/v, 25-50 ml) by sonication (if sonification did not yield a homogenous suspension, a few ml of THF was used to dissolve the solid **S2**, which was then added to the mixture of aqueous H_2_SO_4_). NaNO_2_ (3.86 g, 56.0 mmol) dissolved in H_2_O (2.5 ml) was added slowly to the reaction flask and stirred at room temperature for 1.5 h. In a separate flask H_2_SO_4_:H_2_O (4:3, v/v, 75 ml) was added and heated to reflux. To the refluxing mixture, the **S2** solution was added dropwise and stirred for 30 min. The mixture was quenched with ice water and extracted with EtOAc (3 x 75 ml). After drying the organic layer with Mg_2_SO_4_, the solvent was removed *in vacuo* and the crude product was purified by silica gel flash chromatography (Hex:EtOAc, 2:1) to give **S3**(3.11 g, 50%) as a yellow oil.

^1^H NMR (500 MHz, CDCl_3_) δ ppm 1.249 (t, *J* = 7.5 Hz, 3H), 2.842 (q, *J* = 8.5 Hz, 2H), 7.030 (dd, *J* = 8.5, 2.5 Hz, 1H), 7.230 (d, *J* = 8.5 Hz, 1H), 7.383 (d, *J* = 2.5 Hz, 1H)

^13^C NMR (400 MHz, CDCl_3_) 154.2470, 149.5150, 132.3914, 131.3014, 120.7326, 111.4436, 25.6617, 15.1987

HRMS *(m/z):* [M - H]^−^ calcd for [C_8_H_8_NO_3_]^−^ 166.0510, found 166.0524.

#### tert-butyl 2-(4-ethyl-3-nitrophenoxy)acetate (S4)

Compound **S3** (3.11 g, 18.6 mmol) and tert-butyl 2-bromoacetate (4.35 g, 22.3 mmol) were dissolved in DMF (25 ml). Solid K_2_CO_3_ (5.14 g, 37.2 mmol) was added to the reaction flask and left to stir at +70 °C for 1.5 h until TLC analysis (2:1 Hex:EtOAc, v/v) indicated complete conversion to the product. The solvent was removed *in vacuo* and redissolved in 100 ml EtOAc. The organic layer was washed with saturated NH_4_Cl (50 ml) and brine, then dried over Na_2_SO_4_. Solvent removal *in vacuo* afforded **S4**(4.97 g, 95%) as a yellow oil.

^1^H NMR (500 MHz, CDCl_3_) δ ppm 1.253 (t, *J* = 7.5 Hz, 3H), 1.5 (s, 9H), 2.857 (q, *J* = 7.5 Hz, 2H), 4.554 (s, 2H), 7.116 (dd, *J* = 8.5, 2.5 Hz, 1H), 7.277 (d, *J* = 8.5 Hz, 1H), 7.391 (d, *J* = 3 Hz, 1H)

^13^C NMR (400 MHz, CDCl_3_) 167.3932, 156.3652, 149.4567, 132.3149, 132.2128, 120.6270, 110.0036, 83.0951, 66.0518, 28.1809, 25.7638, 15.1222

HRMS *(m/z):* [M + Na]^+^ calcd for [C_14_H_19_NO_5_Na]^+^ 304.1155, found 304.1160.

#### tert-butyl 2-(4-(1-bromoethyl)-3-nitrophenoxy)acetate (S5)

Compound **S4** (4.97 g, 17.7 mmol), N-bromosuccinimide (3.8 g, 19.5 mmol) and benzoylperoxide (0.2 g, 1 mmol) were dissolved in CCl_4_ (100 ml) and refluxed for 4 h. The reaction mixture was cooled to room temperature and washed with 0.1% NaHCO_3_ (aq) and brine, then dried over Na_2_SO_4_. The solvent was removed *in vacuo* and the crude product was purified by silica gel flash column chromatography (3:1 Hex:EtOAc, v/v) to afford **S4**(5.7 g, 90%) as a yellow oil.

^1^H NMR (500 MHz, CDCl_3_) δ ppm 1.498 (s, 9H), 2.054 (d, *J* = 7 Hz, 3H), 4.571 (s, 2H), 5.787 (q, *J* = 7 Hz, 1H), 7.184 (dd, *J* = 8.5, 3 Hz, 1H), 7.299 (d, *J* = 2.5 Hz, 1H), 7.784 (d, *J* = 9 Hz, 1H)

^13^C NMR (400 MHz, CDCl_3_) 167.0028, 157.7588, 148.0010, 131.1486, 130.8123, 120.7031, 109.7326, 83.3722, 66.0153, 42.0634, 28.1845, 27.3715

HRMS *(m/z):* [M - Br]^+^ calcd for [C_14_H_18_NO_5_]^+^ 280.1179, found 280.1163.

#### 2-(4-(1-bromoethyl)-3-nitrophenoxy)ethan-1-ol (S6)

Compound **S5**(5.7 g, 15.9 mmol) was dissolved in 100 ml THF and cooled down to −78 °C. DIBAL-H (39.8 mmol) was added to the reaction flask and stirred at −78 °C for 20 min, and then left to stir for an additional 2 h at 0 °C. TLC analysis (3:1 Hex:EtOAc, v/v) indicated essentially complete conversion to the product. The reaction was quenched by slowly adding 30 ml H_2_O to the mixture, followed by the addition of 5% HCl (aq) solution until the aqueous solution became acidic (pH = ~4, as judged by pH paper). After vigorously mixing the biphasic mixture in a separatory funnel, the separated organic layer was washed with brine, then dried over Na_2_SO_4_. The solvent was removed *in vacuo* and the crude product was purified by silica gel flash column chromatography to yield **S6**(3.23 g, 60%) as a yellow oil.

^1^H NMR (500 MHz, CDCl_3_) δ ppm 2.056 (d, *J* = 5 Hz, 3H), 4.006 (dd, *J* = 4.5, 4.5 Hz, 2H), 4.142 (dd, *J* = 4, 4 Hz, 2H), 5.785 (q, *J* = 7 Hz, 1H), 7.201 (dd, *J* = 8.5, 2.5 Hz, 1H), 7.356 (d, *J* = 2.5 Hz, 1H), 7.783 (d, *J* = 8.5 Hz, 1H)

^13^C NMR (400 MHz, CDCl_3_) 158.5634, 148.1515, 131.0462, 130.2806, 120.5139, 109.6535, 70.1749, 61.2504, 42.1290, 27.3423

HRMS *(m/z):* [M - Br]^+^ calcd for [C_14_H_18_NO_5_]^+^ 210.0761, found 210.0761.

#### 1-(4-(2-hydroxyethoxy)-2-nitrophenyl)ethan-1-ol (S7)

**S6** (3.23 g, 11.1 mmol) was dissolved in 250 ml H_2_O and refluxed for 1 h. TLC analysis (1:1 Hex:EtOAc, v/v) indicated essentially complete conversion to the product. The product was extracted with EtOAc (3 × 50 ml). The organic layer was washed with brine and then dried over Na_2_SO_4_. The solvent was evaporated *in vacuo* and **S7**(2.0 g, 80%) was used for the next step without further purification.

^1^H NMR (500 MHz, CDCl_3_) δ ppm 1.540 (d, *J* = 6.4 Hz, 3H), 3.997 (dd, *J* = 4.6, 4.6 Hz, 2H), 4.120 (dd, *J* = 7.1, 7.1 Hz, 2H), 5.341 (q, *J* = 6.2 Hz, 1H), 7.201 (dd, *J* = 8.8, 2.8 Hz, 1H), 7.410 (d, *J* = 2.7 Hz, 1H), 7.734 (d, *J* = 8.8 Hz, 1H)

#### 1-(4-(2-(acryloyloxy)ethoxy)-2-nitrophenyl)ethyl acrylate (o-NBMA, S8)

To a solution of **S7**(2.0 g, 8.88 mmol) and acryloyl chloride (26.6 mmol) in CH_2_Cl_2_ (75 ml), TEA (3.5 eq) was added and the mixture was stirred at room temperature for 24 h. The mixture was washed with H_2_O and brine and then dried over Na_2_SO_4_. The solvent was evaporated *in vacuo* and the crude material was purified by silica gel flash column chromatography (2.5:1 Hex:EtOAc, v/v) to yield **S8**(1.79 g, 60%) as a yellow oil.

^1^H NMR (500 MHz, CDCl_3_) δ ppm 1.653 (d, *J* = 6.5 Hz, 3H), 4.253-4.272 (m, 2H), 4.517-4.536 (m, 2H), 5.849 (dd, *J* = 16.5, 1.5 Hz, 1H), 5.87 (dd, *J* = 16.5, 1.5 Hz, 1H), 6.135 (dd, *J* = 33, 10.5 Hz, 1H), 6.135 (dd, *J* = 10.5, 1.5 Hz, 1H), 6.333 (dd, *J* = 6.5, 6.5 Hz, 1H), 6.425 (dd, *J* = 38.5, 1.5 Hz, 1H), 6.425 (dd, *J* = 4, 1 Hz, 1H), 7.181 (dd, *J* = 8.5, 2.5 Hz, 1H), 7.471 (d, *J* = 2.5 Hz, 1H), 7.547 (d, *J* = 8.5 Hz, 1H)

^13^C NMR (400 MHz, DMSO-d6) 165.3601, 164.6091, 157.8246, 148.5720, 132.2687, 132.0463, 128.7689, 128.2840, 127.9741, 127.9522, 120.6063, 109.4215, 67.2670, 66.6545, 62.5167, 21.2189

HRMS *(m/z):* [M + Na]^+^ calcd for [C_16_H_17_NO_7_Na]^+^ 358.0897, found 358.0888.

### Fabrication of photoresponsive polyacrylamide hydrogels

Photoresponsive polyacrylamide gel substrates were prepared based on a previously reported method (*44*). Briefly, Grid-500 high glass-bottom dishes (Fischer, 50-305-810) were activated for gel attachment by sequential treatment with 0.1 M NaOH, 97% (3-aminoproyl)trimethoxysilane (Sigma Aldrich, 281778) and 0.5% glutaraldehyde (Polysciences, 01909). A prepolymer mixture of 40% (w/v) acrylamide solution (25% by volume, Fisher, BP1402), 2% (w/v) bis-acrylamide solution (2.5% by volume, Fisher, BP1404), 50 mM o-nitrobenzyl bis-acrylate (in DMSO, 3.25% by volume), 1M HEPES (pH 7.0, 1% by volume, Sigma Aldrich, H6147) solution, 71.7 mM acrylic acid N-hydroxysuccinimide ester (in DMSO, 4% by volume, Sigma Aldrich, A8060), and H2O (63.25% by volume) was prepared. After degassing for 30 min, polymerization was initiated by adding 10% (w/v) APS (0.6% by volume, Bio-Rad, 161-0700) solution and TEMED (0.4% by volume, Fisher, BP150). Immediately after initiation, 200 μL of gel solution was pipetted onto the activated glass culture dish and covered with a fibronectin-patterned glass coverslip face down (fabricated as described below). After 30 min of polymerization, PBS was added on the dish and the coverslip was removed. Finally, the gel was washed with PBS.

### Preparation of 1D fibronectin micropatterns

1D lines of fibronectin were created on the photoresponsive hydrogels following a microcontact printing method widely applied in the field of surface protein fabrication (*45*). Briefly, PDMS stamps fabricated by photolithography and containing topographical patterns (21 μm width, 40 μm spacing) were obtained from the M. Piel laboratory (Inst. Curie) and used as received (*46*). The patterned side of the stamp was inked with 100 μg/ml fibronectin (Sigma Aldrich, F1141) for 1 h. After drying the stamp using a stream of air, the fibronectin-coated stamp was stamped onto a 12 mm no. 1.5 circular coverslip (Fisher, 12-545-80), rinsed with ethanol and treated with plasma (Harrick Plasma) for 60 sec, and a 20 g weight was placed on top of the stamp. The fibronectin pattern was finally transferred to the gel surface by placing the coverslip face down on the prepolymer solution as described above, immediately upon the initiation of polymerization.

### Fabrication of stepwise stiffness gradients by controlled UV exposure

Stiffness patterns were fabricated on photoresponsive hydrogels using a Nikon Eclipse Ti-E epifluorescence microscope and Plan Fluor 10x/0.30NA objective (Nikon), controlled by NIS-Elements software (Nikon). The fibronectin-patterned photoresponsive gel was placed on the stage and, using phase-contrast imaging, two regions were selected such that they were ‘A’ mm (A > 2) apart. A hypothetical line connecting the two regions ran across the fibronectin patterns perpendicularly (Fig. S3). The field diaphragm lever was then adjusted so that the diameter of the illuminated area on the substrate was 500 μm. Fluorescence imaging using a 395/25 nm LED (315 mW) and DAPI filter set with LED fluorescence illumination from a SpectraX light Engine (Lumencor) was initiated, and a time lapse movie of the two regions was captured at 0 s intervals for ‘15 × A’ min, leaving the active shutter open during stage movement. This led to a 500 μm x ‘A’ mm region being photoirradiated to the extent that all the photolabile crosslinkers in the exposed region were cleaved. The process was repeated in regions parallel to and 500 μm apart from the first irradiated area, resulting in a gel that had alternating, 500 μm wide stiff (~15 kPa) and soft (~8 kPa) regions.

### Stiffness characterization by bead indentation

The irradiation time-dependent change in the Young’s modulus of the photoresponsive polyacrylamide gel was measured using a bead indentation method (*17*) based on Hertzian indentation theory. A thick (>1 mm) hydrogel was created by pipetting 300 μl of prepolymer solution onto an activated glass culture dish and covering it with a 25 mm no. 1.5 circular coverslip (Fisher, 12-545-102). After polymerization, the coverslip was removed in PBS and the gel was washed with additional PBS. A silica bead (Polysciences, 1 mm diameter) was placed on the gel after 200 nm crimson fluorospheres were first gravity-settled on the gel surface to function as markers for measuring bead contact area with epifluorescence microscopy. At each irradiation time point, the bead indentation depth (*δ*) was calculated from the bead radius (*R*) and the contact radius (*r*) according to equation (1):

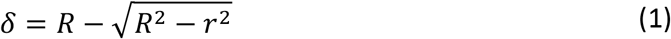

From this indentation depth, the Young’s modulus (*E*) was calculated using the Poisson ratio of the hydrogel (*v*) and buoyancy corrected bead force (*f*) according to the Hertz solution:

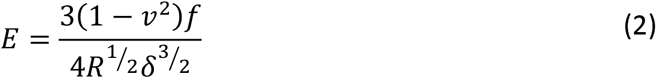

For polyacrylamide gels, *v* = 0.3‒0.5 (here, *v* = 0.3 was used). The glass bead density was measured to be ~2600 kg/m^3^.

### Immunofluorescence staining

Samples were fixed for 10 min with warm 4% PFA, followed by permeabilization and blocking for 20 min with 0.3% Triton X-100 in 10% horse serum (Gibco, 16050-122). Primary antibodies were diluted in 10% horse serum and samples were incubated with the antibody overnight at +4 °C. Secondary antibodies were diluted in PBS and samples were incubated with the antibody for one to two hours at room temperature. Where indicated, the nuclei were counterstained using 5 μg/ml DAPI (4’,6-diamidino-2-phenylindole) or 500 nM SiR-DNA (Spirochrome, SC007; for live cells) and filamentous actin using 200 nM SiR-actin (Spirochrome, SC001).

### Fluorescence and brightfield microscopy

Most fluorescent specimens were imaged using a Marianas spinning disk confocal microscope with a Yokogawa CSU-W1 scanning unit, controlled by SlideBook 6 software (Intelligent Imaging Innovations). The objectives used were a 20x/0.8 Plan-Apochromat (Zeiss) and 40x/1.1 W LD C-Apochromat (Zeiss), and images were acquired using an Orca Flash4 sCMOS camera (Hamamatsu Photonics). The 2D gradient hydrogels with cells were imaged using Nikon Eclipse Ti2-E widefield microscope, controlled by NIS-Elements AR 5.11 software (Nikon). The objective used was a 10x/0.3 CFI Plan-Fluor objective (Nikon), and images were acquired using an Orca Flash4 sCMOS camera (Hamamatsu Photonics) and 2×2 binning.

Live phase contrast imaging of U-251MG cells on photoresponsive hydrogels was done using a Nikon Eclipse Ti-E microscope and an Andor Zyla 5.5 sCMOS camera (Andor Technology). The objective was a Plan Fluor 10x/0.30NA objective (Nikon) and samples were maintained in a Bold Line stage top humidified incubator (Okolab) at +37 °C/5% CO_2_.

### Traction force microscopy

To measure the tractions exerted by MDA-MB-231 cells on their substrate, polyacrylamide hydrogels of varying stiffness (fibronectin-functionalized and supplemented with fluorescent microbeads) were manufactured on glass-bottom dishes as described above. Cells were seeded on the gels (5,000 cells/plate) approximately 24 hours after transfection with the indicated siRNAs, and grown for another 48 hours before the experiment was conducted. For imaging the cells and beads, a Marianas spinning disk confocal microscope with a stage top incubator unit (+37 °C/5% CO_2_) was used. Brightfield images of single cells and fluorescence z-stacks of the beads embedded in the hydrogel were captured before and after cell detachment by addition of 2% sodium dodecyl sulfate.

The resulting data were analyzed using a previously described implementation of Fourier transform traction cytometry (*47*). First, displacement fields were calculated using high-resolution subsampling and assuming no outward deformation of the substrate. Optimal L2-regularization was performed on sets of images acquired from soft and stiff gels to determine the final regularization parameter λ = 5 × 10^−6^, which was then used for calculating all the subsequent traction fields. The background, or noise, of the measurements was estimated by analyzing five empty (i.e. no cells) fields of view per substrate stiffness.

### Finite element analysis

To estimate the effective spring constant around the interface of a stepwise stiffness gradient, a finite element model using COMSOL Multiphysics (COMSOL, Inc., Burlington, MA) multibody dynamics module was utilized. Two three-dimensional blocks (120 μm × 60 μm × 20 μm) were created and interfaced at (x = 0). Linear elastic material properties were prescribed to both blocks with Poisson’s ratio = 0.4, density = 1000 kg/m^3^ and Young’s modulus = 1 kPa and 10 kPa. A lateral 0.5 nN force was applied on a circular (1 μm radius) surface contact (Figure S5a). Fixed boundary conditions were applied to all surfaces except the top surface. The displacement field due to applied loads was computed on a model created using built-in automatic meshing routines (extra-fine mechanics-based mesh). These data were used to calculate effective spring constant at the contact zone (*K*_*eff*_ = applied force/average displacement under the circular contact area). The location of the circular contact and direction of the force were varied, and effective spring constants were calculated accordingly (Figure S5b).

### Computational modeling of single-cell migration and growth cone steering on stiffness gradients

A previously described (*37*) C++ version of the stochastic cell migration simulator (CMS) was modified to account for spatial variations in substrate stiffness. The detailed algorithms and equations governing the base CMS have been described in full in (*28*). Briefly, the CMS uses the Gillespie Stochastic Simulation Algorithm (*48*) to simulate an entire cell by connecting several motor-clutch modules to a central cell body and then balancing forces at the center (Fig. 3a). Here, the cells were simulated for 60 min to allow them to reach a dynamic steady state, after which each cell was displaced randomly to a 180 μm × 180 μm region on a substrate with repeating soft and stiff areas and connecting stiffness gradients (Fig. 3b). Cell positions and traction forces were recorded every second and used to calculate RMC and mean traction force per module. Custom Matlab code was used to quantify module forces on soft and stiff substrates, and to track the displacement of individual cells, from gradients or soft regions, over time. All the CMS simulations were conducted at the Minnesota Supercomputing Institute (MSI). For additional details on the cellular level model and its implementation, see Supplementary Text 2.

The CMS was further modified to investigate filopodial and GC dynamics on substrate stiffness gradients. The filopodia were represented by individual CMS modules that were arranged around an initially semicircular GC. Each filopodia was allocated a set number of molecular clutches – the corresponding substrate clutches were distributed randomly and their spring constants varied linearly with position along the gradient. The details of the GC model and corresponding simulations are presented in Supplementary Text 3.

### Image analysis

Images were analyzed using ImageJ (National Institutes of Health) and CellProfiler v2.2.0 (Broad Institute) software. For analysis of YAP nuclear localization, a custom CellProfiler pipeline was used to segment the cells into nuclei (corresponding to the nuclear counterstain) and cytoplasm (a region of max. 4 μm around the nucleus, excluding parts outside the cell). The mean gray value in the nucleus was divided by the corresponding value in the cytoplasm. For analysis of vinculin-positive adhesions in MDA-MB-231s, a semi-automatic ImageJ script was used: an individual confocal plane from the basal side of the cell was subjected to background removal (rolling ball) and thresholding to exclude cytoplasmic signal and peripheral ruffles. The number and sizes of the remaining adhesions were recorded.

### Statistical analysis

Statistical analyses and plotting were performed using GraphPad Prism v6.05 (GraphPad) and R v3.5.1 (R Core Team). Confidence intervals for means were calculated using bias-corrected and accelerated (BCa) bootstrap intervals from 10,000 resamples. Confidence intervals for binomial data were calculated using Wilson score interval. Whenever data were deemed to follow a non-normal distribution (according to Shapiro-Wilk normality test), analyses were conducted using non-parametric methods. The names and/or numbers of individual statistical tests, samples and data points are indicated in figure legends. Unless otherwise noted, all results are representative of three independent experiments and two-tailed *p*-values have been reported.

## Data Availability

The data supporting the findings of this study are available within the article and from the authors on reasonable request. All code and scripts are available online (oddelab.umn.edu and GitHub, https://github.com/cbcbcbcb123/Growth-Cone-Dynamics) or on request from the corresponding authors.

## Conflicts of Interest

The authors declare no competing interests.

## Acknowledgements

We thank Louis S. Prahl, Jin Tian and Guoyou Huang for helpful discussions on computational modeling and the Ivaska lab members for their insightful comments and discussion. Simulations were run in part on high-performance computing resources at the Minnesota Supercomputing Institute. Turku Bioscience Centre Cell Imaging Core and Biocenter Finland are acknowledged for services, instrumentation and expertise. The authors are supported by the University of Turku Doctoral Programme in Molecular Life Sciences (A.I), the Academy of Finland (312517, J.I.), ERC CoG grant (615258, J.I.), Sigrid Juselius Foundation (J.I.), the Finnish Cancer Organization (J.I.), the National Natural Science Foundation of China (11972280, F.X.; 11772253, M.L.; 12022206, M.L.; 11532009, T.J.L), the Shaanxi Province Youth Talent Support Program (M.L.), the Young Talent Support Plan of Xi’an Jiaotong University (M.L.), the National Institutes of Health (R01 AR077793, G.M.G.; R01 CA172986, D.J.O.; U54 CA210190, D.J.O.), and the NSF Science and Technology Center for Engineering Mechanobiology (CMMI 1548571, G.M.G).

## Author Contributions

Conceptualization, A.I., K.-Y.P., J.H., B.C., B.F., J.K., M.L., M.D.D., J.I., D.J.O.; Formal Analysis, A.I., K.-Y.P., J.H., B.C.; Funding Acquisition, M.L., T.J.L., G.M.G., F.X., M.D.D., J.I., D.J.O.; Investigation, A.I., K.-Y.P., J.H., B.C., G.S., B.F., J.K., M.M.M., F.X.; Methodology, A.I., K.-Y.P., J.H., B.F., J.K., M.L., F.X., M.M.M., T.J.L., G.M.G.; Project Administration, M.L., T.J.L., F.X., M.D.D., J.I., D.J.O.; Resources, M.L., F.X., T.J.L., G.M.G., M.D.D., J.I., D.J.O.; Software, A.I., J.H., B.C., F.X.; Supervision, M.L., F.X., T.J.L., G.M.G., M.D.D., J.I., D.J.O.; Validation, A.I., K.-Y.P., J.H., B.C., M.L., G.M.G.; Visualization, A.I., K.-Y.P., J.H., B.C., G.S., M.L., F.X., T.J.L., G.M.G.; Writing – Original Draft, A.I., K.-Y.P., B.C., M.L., G.M.G., M.D.D., J.I., D.J.O.; Writing – Review & Editing, all authors

---

**Supplementary Text 1: Chemistry of o-NBbA and photoresponsive polyacrylamide hydrogels**

Polyacrylamide was selected as the base material for the stiffness gradients used in this study, as it is the most widely employed model system for investigating the role of substrate stiffness in directing cell behavior. This is partly due to the ease of obtaining elastic moduli in a wide, physiologically relevant range (*49*, *50*). While other types of gels (e.g. collagen or hyaluronic acid) are known to interact directly with cell surface receptors, including integrins, polyacrylamide gels are inert to such interactions. This allows more control over the types and densities of ligands that will be presented to the cells, making the material ideal for mechanobiological studies. Various methods have been developed to fabricate stiffness gradients in gels to study the durotactic behavior of cells. Some examples exploit the diffusion of two prepolymer solutions (*12*, *30*), tilted-superposition of two hydrogels (*51*), freeze-thaw-induced crosslinking of polyvinyl alcohol (*52*), or toehold-mediated strand displacement of DNA (*53*). Aiming for high-resolution spatiotemporal control over the mechanical properties of the gel, we chose light as the external stimulus (*54*–*58*). Therefore, we aimed to design and synthesize a new, minimalistic and photocleavable crosslinker that contains acrylate moieties.

Among various photolabile functionalities that are available, o-nitrobenzyl (o-NB) was chosen due to its high one-photon photolysis efficiency and high deprotection yields (*59*, *60*). o-NB based compounds have been used widely in hydrogel-based studies to achieve controlled release or immobilization of payloads (*61*–*65*), photodegradation of gels (*66*–*68*), or modulation of gel stiffness (*56*, *69*). However, many of these studies have focused on polyethylene glycol-based gels rather than polyacrylamide, or complete degradation of the gel rather than controlling the Young’s modulus. To our knowledge, there has been only one report to date where o-NB-based crosslinkers have been used to fabricate photoresponsive polyacrylamide hydrogels (*70*). While the study demonstrates the feasibility of o-NB based crosslinking, the method itself requires multiple steps to crosslink chains of polyacrylamide through the o-NB moiety, which made its application here unwieldy.

In this study, a simple one-step synthesis of photoresponsive polyacrylamide gels was enabled by the functionalization of an o-NB group with two acrylate moieties to yield a crosslinker that would cleave upon photolysis. The photocleavable crosslinker, o-nitrobenzyl bis-acrylate (o-NBbA), was synthesized in seven steps from p-ethyl aniline (Fig. S2a) and designed so that its cleavage would not release any byproducts in the medium (Fig. S2b). Based on a previously reported polyacrylamide recipe (*44*), a photoresponsive gel with an initial stiffness of 20 kPa that can be reduced down to 10 kPa was designed by replacing 50 mol % of bis-acrylamide with o-NBbA. The resulting gel exhibited a Young’s modulus of ~15 kPa that was reduced down to ~8 kPa after complete cleavage of the o-NBbA crosslinker by exposure to 395 nm light for 5 min (Fig. S3b–d). The slight discrepancy between the expected and measured substrate stiffness could be due to the relatively low water solubility and partial phase separation of o-NBbA in the prepolymer solution, which would result in softening of the hydrogel post-polymerization (*71*).

The light source used for the photocleavage was an LED from a SpectraX light engine instrument installed in a Nikon TiE microscope, originally intended for epifluorescence imaging. This method had several advantages: spatial control can be achieved easily, as the location of the substrate can be precisely chosen via phase-contrast imaging and the area of irradiation can be controlled with the field diaphragm and objectives. For instance, the diameter of the LED-irradiated area could be adjusted to as low as 59 μm using a 40x objective with a nearly closed field diaphragm, or as high as 978 μm under a 10x objective with a fully opened field diaphragm. Stiffness patterns could also be created using the ‘time lapse movie’ function of the NIS-Elements software (Nikon). Here, alternating stiffness gradients were created by initiating a time-lapse movie between two regions of the gel, a method that could be modified to yield more complex 1D patterns or even 2D shapes. Although not explored here, temporal control would be equally possible: for example, stiffness gradients could be introduced in gels at various time points during live cell culture, while simultaneously observing cellular behavior and responses.

To conjugate fibronectin to the surface of the gel via covalent interaction, acrylic acid N-hydroxysuccinimide (NHS) ester was used as the tethering agent. While two methods, addition of acrylic acid NHS ester in the pregel solution followed by stamping of fibronectin, or stamping of the pregel solution with fibronectin preincubated with acrylic acid NHS ester, both produced fibronectin-patterned hydrogels, the former was chosen since it yielded more consistent results. Once the gel had been fabricated, any remaining NHS ester moieties in the gel were passivated with bovine serum albumin (BSA) in PBS to prevent any non-specific interactions between the gel and cells.

**Supplementary Text 2: Implementation of the cell migration simulator using mechanically heterogeneous substrates**

To establish whether our observations of negative (and positive) durotaxis could be explained through a single set of principles, namely the motor-clutch dynamics of cell-matrix adhesions, we developed a modified version of the cell migration simulator (CMS) that can be used for modeling cell migration on mechanically heterogeneous substrates. The detailed governing equations and algorithms of the original CMS were described previously (*28*). Briefly, the CMS comprises multiple motor-clutch models (i.e. modules) that mimic cellular protrusions, and determines cell motion by a force balance between the modules and a central cell body (Fig. 3a). In the CMS, new modules are nucleated stochastically, module length increases over time via actin polymerization that is simultaneously counteracted by myosin-induced retraction of actin fibers, and modules are removed when they become too short. In addition, total actin and numbers of clutches and motors are kept constant in accordance with the conservation of mass.

In each motor-clutch system, adhesion clutches bind to elastic substrate springs with a constant rate of *K*_*on*_. Connected clutches form a direct mechanical link from the intracellular cytoskeleton to the extracellular substrate – forces are borne from active myosin motors and transmitted by the resulting inward actin flow. The unbinding rate of a connected clutch *i*, *K*_*off,i*_, varies with force *F*_*i*_ according to the Bell model (*72*):

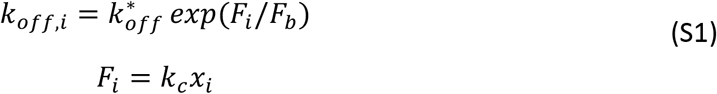

where 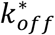 is the clutch unbinding rate in the absence of loading, *F*_*b*_ is the characteristic clutch rupture force, and *x*_*i*_ is the elongation of the spring representing the *i*^*th*^ connected clutch with a spring constant *K*_*c*_. The actin filaments are pulled by *n*_*m*_ myosin motors, each capable of exerting a force *F*_*m*_, and balanced by the traction force *F*_*s*_, resulting in inward actin flow with the effective actin flow rate (*v*_*m*_) based on

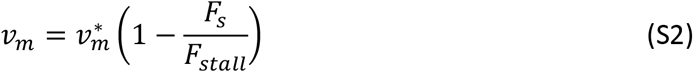

where 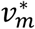 is the unloaded rate, *F*_*stall*_ = *n*_*m*_*F*_*m*_ is the stall force of the ensemble of myosin motors, and the traction force *F*_*s*_ transmitted by all the connected clutches is given by:

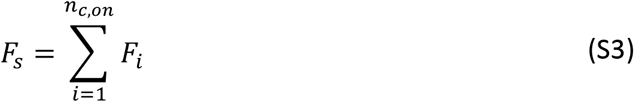

in which *n*_*c*,*on*_ is the number of connected clutch bonds. Actin monomers are added to the barbed ends of actin filaments in the cellular protrusions (modules) at a polymerization rate *v*_*p*_, constrained by the total actin length *A*_*tot*_ in the cell according to the relation:

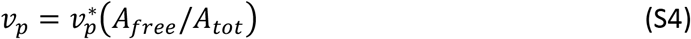

where *A*_*free*_ is the amount of available G-actin and 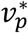 is the maximum polymerization rate. Module elongation and retraction both result from this actin polymerization and the actin flow rate (*v*_*m*_). New modules are nucleated at a nucleation rate *K*_*mod*_, also constrained by actin availability:

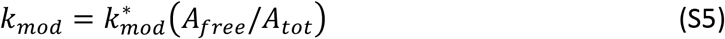

where 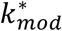 is the maximum module nucleation rate. Actin filaments are depolymerized into actin monomers when they pass through the position of the myosin motors. Filaments can also be capped and polymerization arrested by actin capping proteins at a capping rate *K*_*cap*_. Actin filaments, and the corresponding modules, are removed from the simulation when their length falls below *l*_*min*_.

Monte Carlo simulations were conducted using a direct Gillespie Stochastic Simulation Algorithm (*48*), with each time step determined based on total event rates, including *K*_*on*_, *K*_*off,i*_, *K*_*mod*_, and *K*_*cap*_, and the event execution determined based on accumulated event rates. The CMS C++ version, described in (*37*), was modified to account for variations in substrate stiffness (described below), and simulations were conducted in Mesabi computer cluster at the Minnesota Supercomputing Institute (MSI).

After the simulated cells had reached a dynamic steady state (60 min), they were displaced randomly to a 180 μm × 180 μm region (Fig. 3b), and the substrate stiffnesses (*K*_*s*_) experienced by the cell body and each protrusion were determined based on their respective y-coordinates (*y*). The substrate could be either soft (*K*_*soft*_), stiff (*K*_*stiff*_), or between the two extremes [gradients following a normal cumulative distribution function, described by the following error functions (erf)]:

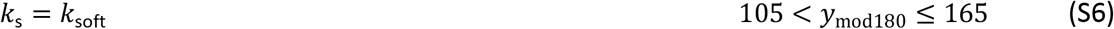

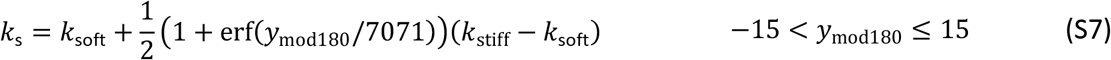

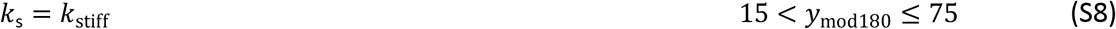

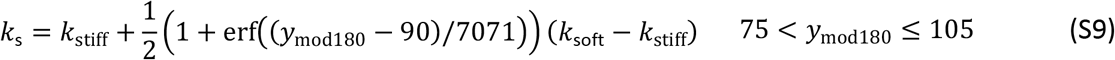

where *y*_mod180_ = (*y* mod 180)/*y*_0_ with (*y* mod 180) representing the remainder of *y* divided by 180 μm (ranging from −15 to 165) and *y*_0_ = 1 μm. This way, the number of cells in both soft and stiff regions was initially the same. In addition, by repeating the same stiffness pattern *ad infinitum*, the finite amount of cells placed in the finite rectangular region was representative of infinite cells placed on an infinite substrate with the same initial distribution of cells between soft and stiff areas. A normal cumulative distribution function was selected due to a finite element model of polyacrylamide, which demonstrated that the effective spring constant around a stepwise gradient of elastic modulus follows a similar distribution (Fig. S5). This was true regardless of the orientation of the applied traction (soft-to-stiff vs. stiff-to-soft).

Here, we adopted the high-motor-clutch parameter values used in (*28*) to describe U-251MG migration on mechanically distinct but isotropic substrates (Table S2). Clutch stiffness was further adjusted to 8 pN nm^−1^ to better recapitulate the stiffness-dependence of U-251MG speed *in vitro* (*28*). During the CMS simulations, cell positions and traction forces were recorded every second. The data collected during the first 60 min were analyzed to ensure that the simulated cells had indeed reached a dynamic steady state. Random motility coefficients (RMC) were calculated as described previously (*28*). Briefly, the mean squared displacement, 〈*r*^2^〉, was calculated with overlapping time periods Δ*t* = 10 min, 20 min,&, and plotted as a function of Δ*t*. The first half of the plotted curve was fitted with a straight line (slope = 〈*r*^2^〉/Δ*t*), and RMC was given by RMC = 〈*r*^2^〉/4Δ*t*. Module forces were recorded every 10 min and averaged throughout the simulation to yield the average traction force per module. Custom Matlab scripts were employed to analyze the change in cell numbers in soft and stiff regions over time, to compare module forces in the soft and stiff parts of the gradients, and to track individual cells over time based on their initial location in soft or graded substrate regions.

On 10‒100 pN nm^−1^ gradients, we found that the majority of cells translocated away from stiffer regions and toward soft areas (Fig. 3e‒f), which were associated with higher traction forces per module and lower overall migration speed, RMC (Fig. 3c‒d). We also tested whether altering the range of the gradient would affect the durotaxis. On 0.3‒3 pN nm^−1^ gradients, the stiffer side was associated with higher traction forces and higher RMC (Fig. S6a‒b). On these substrates, simulated cells displayed rapid accumulation in the stiffer regions (Fig. S6c). Finally, when the gradient was chosen such that there would be no appreciable difference in mean traction forces (100‒300 pN nm^−1^), cells clustered primarily in stiffer regions with lower RMC (Fig. S6a‒b,d).

**Supplementary Text 3: Modeling axonal pathfinding and mechanosensitive steering of growth cones**

Axonal growth cones (GCs) (Fig. S7a) can turn or contract in response to substrate stiffness gradients (*23*, *73*) by controlling the dynamics of adhesions, filopodial remodeling, and active contraction (*38*). To establish whether motor-clutch dynamics could explain the mechanosensitive turning of neuronal GCs (*23*), akin to the negative durotaxis exhibited by the U-251MG glioblastoma cells, we modified the CMS to model an individual GC on a functionally graded substrate. A group of *i* filopodia, each modeled as a single molecular clutch module (Figs. 7b), were attached to a GC central domain. Each module was allocated *n*_*i*_ molecular clutches (linear springs of stiffness *K*_*c*_) and *n*_*i*_ corresponding substrate clutches (linear springs of stiffness *K*_*s,i*_). Substrate clutches were distributed randomly, and had values *K*_*s,1*_ ≤ *K*_*s,i*_ ≤ *K*_*s,2*_ that varied linearly with position along the gradient.

Monte Carlo simulations were conducted to evaluate filopodial and GC dynamics over time. We modeled a GC as having 21 potential growth sites for filopodia, chosen from a uniform orientation distribution between −π/2 and π/2, relative to the direction of the ‘axon’. New protrusions with an initial length *l*_*in*_ and width *l*_*wid*_, dictating the effective clutch-ligand binding area, were added into the simulation at a rate *K*_*mod*_ and assigned *n*_*m*_ myosin motors; note that we used an actin filament in the schematic diagram (Fig. S8a) to represent the filament bundle in the filopodium. The adhesion and substrate clutches under each filament then evolved according to the clutch binding and unbinding dynamics described above. Unlike the cellular level CMS, our modified model assumes a relatively stable pool of actin monomer in the GC. Thus, the actin polymerization rate *v*_*p*_ remained constant during each simulation. See Table S3 for parameter details.

First, we investigated whether the dozens of filopodia within a GC might enable mechanically directed growth by evaluating the response of an individual 8 μm filopodium to a linear stiffness gradient of 0.01 to 100 pN nm^−1^ (Fig. S7b). The filopodium was placed on a 10 μm × 10 μm square substrate and oriented at an angle 0 ≤ θ ≤ π/2 relative to the gradient (Fig. S7c). When the filopodium length was fixed, simply increasing the orientation between the filopodium and the gradient was sufficient to significantly increase traction force generation (Fig. S7d). Conversely, when the orientation was fixed at π/2, i.e. perpendicular to the gradient, we found that both traction force and the number of engaged clutches increased linearly with filopodium length (Fig. S7d).

Next, we investigated the impact of different stiffness gradients for traction force generation using a fixed filopodium length (8 μm) and orientation (0). Maximal traction forces resulted from the filopodium sensing a soft region, in the range of 0.01 to 0.1 pN nm^−1^. The higher end of the stiffness gradient proved significantly less important for the overall traction (Fig. S7e). This result demonstrates that a filopodium can generate comparatively high forces even if only a part of it is located on softer substrate. Thus, high traction force generation by individual filopodia is favored at a low optimal stiffness and forces drastically drop on stiffer matrices.

Higher traction forces are often accompanied by a decrease in actin retrograde flow, as myosin-borne forces are transmitted to the ECM instead of freely displacing actin. Regardless of filopodia orientation, actin in GCs flows toward the structure’s center, and much like traction forces, actin flow rates can also differ for different types of neurons (*39*). We therefore investigated how both the speed and direction of actin flow relative to the stiffness gradient affect the dynamics of single filopodia. By studying filopodia oriented at their growing end with either the stiffer or more compliant end of a stiffness gradient (Fig. S7f), we found that orientation toward the compliant end of the substrate (and hence actin retrograde flow toward the stiff end of the substrate) led to increased extension rates and decreased retraction rates (Fig. S7g). In all cases, the overall growth rate of filopodia was a trade-off between growth at the constant actin polymerization rate *v*_*p*_, and shortening at the actin flow rate *v*_*m*_, which varied almost linearly with substrate stiffness (Fig. S7h). For an intermediate polymerization rate of *v*_*p*_ = 120 nm s^−1^, orientation affected filopodia growth rate by a factor of two (Fig. S7h). Together, these results provide a mechanism by which individual filopodia can exert more traction and elongate faster on softer substrates.

We then investigated whether these changes in filopodial dynamics could contribute to GC steering on stiffness gradients. First, we evaluated the degree of GC turning on one type of stiffness gradient (*K*_*s,1*_ = 0.01 pN nm^−1^ and *K*_*s,2*_ = 100 pN nm^−1^) by studying an initially semicircular GC with 21 uniformly distributed filopodia (Fig. S8a). Within 15 seconds of simulation, the filopodia pointing toward the compliant end of the substrate outgrew the rest, resulting in an effective turning of the GC (Fig. S8a). As expected from the previous results, filopodia in the softer regions of the gradient were longer and generated higher traction forces (Fig. S8b).

To investigate the effect of different stiffness gradients on GC turning in detail, we repeated our simulations over a broad range of possible substrate stiffnesses, with *K*_*s,left*_ and *K*_*s,right*_ varying from 0.01 to 100 pN nm^−1^. To quantify the degree of turning, we defined a parameter 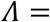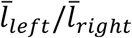, which represents the degree to which the GC has turned left. Here, 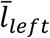 and 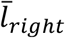 are the average lengths of filopodia in the left-hand and right-hand sides of the GC after 100 seconds of simulation, respectively. In addition to developing polarity through turning, the GC could enlarge, with all filopodia elongating as compared to their initial length, or retract (Fig. S8c). Enlarged GCs appeared on very compliant substrates (red section, *K*_*s,left*_ and *K*_*s,right*_ on the order of 0.01 to 0.1 pN nm^−1^) with a negligible stiffness gradient, and retractile GCs appeared on higher stiffnesses, independent of the actual strength of the gradient (green section, *K*_*s,left*_ and *K*_*s,right*_, on the order of 1 to 100 pN nm-1). Finally, polarized GCs appeared on compliant substrates with a moderate or high stiffness gradient (*K*_*s,left*_ on the order of 0.01 to 0.1 pN nm^−1^, >1 pN/nm/20 μm). A phase diagram for GC turning illustrates how the structure can either remain straight or turn to the more compliant side (Fig. S8d), and reveals that a stronger gradient may also promote GC turning, unless the range of the gradient as a whole is significantly stiffer than the optimal stiffness range for individual filopodia (Fig. S8e). Thus, the motor-clutch model can recapitulate mechanosensitive GC steering toward softer matrix *in silico*.

**Table S1.** Relative acrylamide and bis-acrylamide concentrations and corresponding Young’s moduli for homogeneous (constant modulus) hydrogels.

| Final acrylamide % | Final bis-acrylamide % | Volume of (40%) AA stock, $\mu\text{l}$ | Volume of (2%) bis-AA stock, $\mu\text{l}$ | PBS, $\mu\text{l}$ | ∼Young's modulus, $\text{kPa}^1$ |
| --- | --- | --- | --- | --- | --- |
| 5.4 | 0.04 | 63 | 10 | 397 | 0.5 |
| 5.7 | 0.08 | 63 | 17.5 | 365 | 2 |
| 7.5 | 0.2 | 94 | 50 | 356 | 9.6 |
| 12 | 0.2 | 150 | 50 | 300 | 22 |
| 18 | 0.4 | 225 | 100 | 175 | 60 |

<sup>1</sup>Values obtained using atomic force microscopy, see (30).

**Table S2.** Parameters for the cellular level CMS.

| Parameter | Symbol | Value | Ref. |
| --- | --- | --- | --- |
| Total number of myosin motors | $N_m$ | 10,000 | (28) |
| Total number of clutches | $N_c$ | 7,500 | (28) |
| Maximum total actin length | $A_{tot}$ | 100 $\mu\text{m}$ | (28) |
| Maximum actin polymerization rate | $v_p^*$ | 200 nm/s | (28) |
| Maximum module nucleation rate | $k_{mod}^*$ | 1 $\text{s}^{-1}$ | (28) |
| Module capping rate | $k_{cap}$ | 0.001 $\text{s}^{-1}$ | (28) |
| Initial module length | $l_{in}$ | 5 $\mu\text{m}$ | (28) |
| Minimum module length | $l_{min}$ | 0.1 $\mu\text{m}$ | (28) |
| Cell spring constant | $k_{cell}$ | 10,000 pN/nm | (28) |
| Number of cell body clutches | $n_{c,cell}$ | 10 | (28) |
| Substrate spring constant | $k_s$ | 0.3–300 pN/nm | Adjusted |
| Maximum number of module motors | $n_m^*$ | 1,000 | (28) |
| Myosin motor stall force | $F_m$ | 2 pN | (28) |
| Unloaded actin flow rate | $v_m^*$ | 120 nm/s | (28) |
| Maximum number of module clutches | $n_c^*$ | 750 | (28) |
| Clutch on-rate | $k_{on}$ | 1 $\text{s}^{-1}$ | (28) |
| Unloaded clutch off-rate | $k_{off}^*$ | 0.1 $\text{s}^{-1}$ | (28) |
| Clutch spring constant | $k_c$ | 8 pN/nm | Adjusted |
| Characteristic clutch rupture force | $F_b$ | 2 pN | (28) |

**Table S3.**
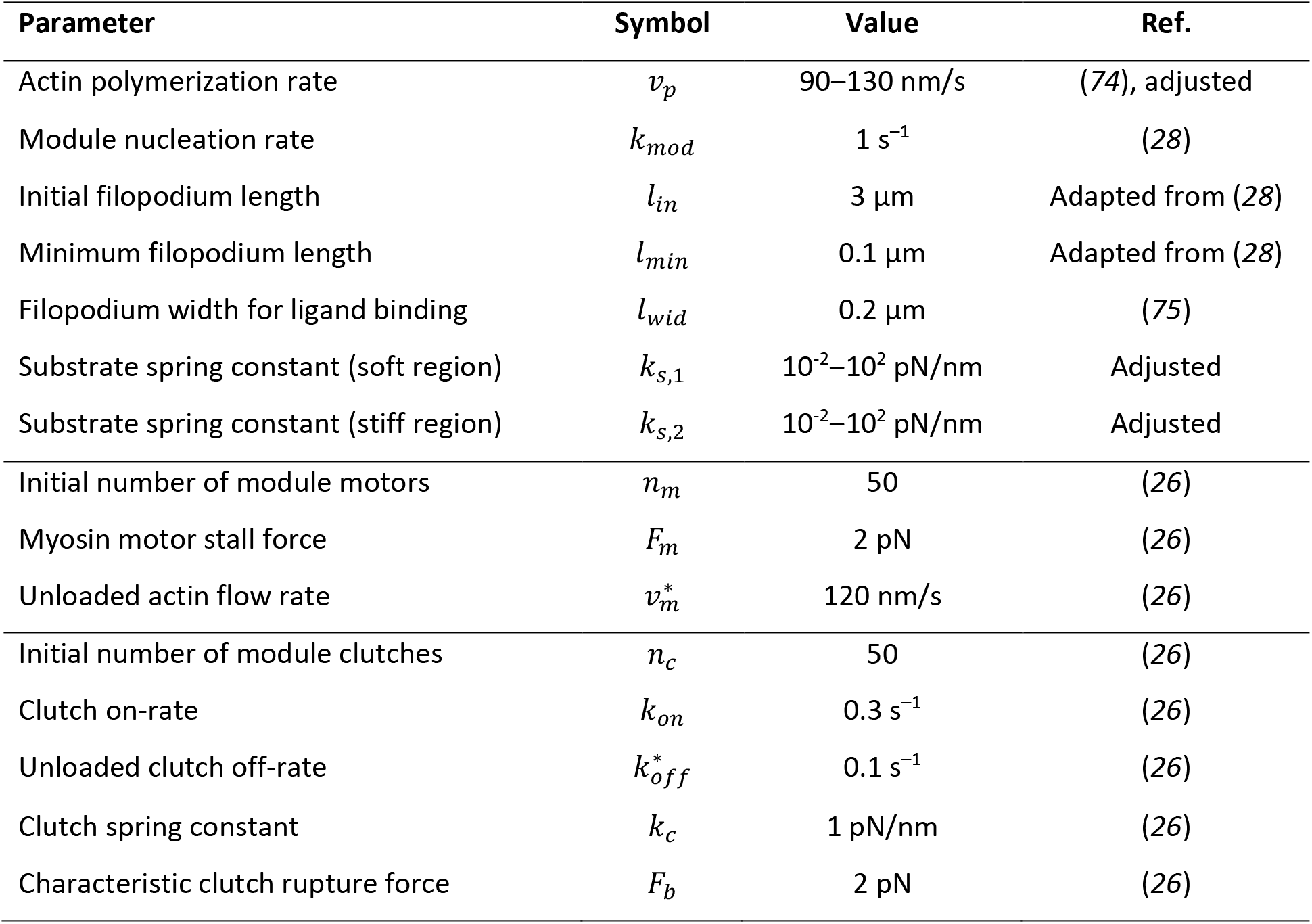
Parameters for the filopodia/GC model.

| Parameter | Symbol | Value | Ref. |
| --- | --- | --- | --- |
| Actin polymerization rate | $v_p$ | 90–130 nm/s | (74), adjusted |
| Module nucleation rate | $k_{mod}$ | $1 \text{ s}^{-1}$ | (28) |
| Initial filopodium length | $l_{in}$ | 3 $\mu\text{m}$ | Adapted from (28) |
| Minimum filopodium length | $l_{min}$ | 0.1 $\mu\text{m}$ | Adapted from (28) |
| Filopodium width for ligand binding | $l_{wid}$ | 0.2 $\mu\text{m}$ | (75) |
| Substrate spring constant (soft region) | $k_{s,1}$ | $10^{-2}$–$10^2$ pN/nm | Adjusted |
| Substrate spring constant (stiff region) | $k_{s,2}$ | $10^{-2}$–$10^2$ pN/nm | Adjusted |
| Initial number of module motors | $n_m$ | 50 | (26) |
| Myosin motor stall force | $F_m$ | 2 pN | (26) |
| Unloaded actin flow rate | $v_m^*$ | 120 nm/s | (26) |
| Initial number of module clutches | $n_c$ | 50 | (26) |
| Clutch on-rate | $k_{on}$ | $0.3 \text{ s}^{-1}$ | (26) |
| Unloaded clutch off-rate | $k_{off}^*$ | $0.1 \text{ s}^{-1}$ | (26) |
| Clutch spring constant | $k_c$ | 1 pN/nm | (26) |
| Characteristic clutch rupture force | $F_b$ | 2 pN | (26) |

## Notes

### Competing Interest Statement

The authors have declared no competing interest.

